# ALK Variants Differentially Modulate Neuroblastoma Tumor Behavior and Transcriptome in a Cellular Context-Dependent Manner

**DOI:** 10.64898/2026.09.08.750106

**Authors:** Maya El Natour, Lucie Vivancos Stalin, Nastassia Gobet, Viviane Praz, Katia Balmas Bourloud, Nicolas Jauquier, Isabelle Janoueix-Lerosey, Nicolò Riggi, Raffaele Renella, Annick Mühlethaler-Mottet

**Affiliations:** Pediatric Hematology-Oncology Research Laboratory, Woman-Mother-Child Department, Lausanne University Hospital and University of Lausanne, Lausanne, Switzerland; Experimental Pathology, Laboratory Department, Lausanne University Hospital and University of Lausanne, Lausanne, Switzerland; Pediatric Surgery, Woman-Mother-Child Department, Lausanne University Hospital and University of Lausanne, Lausanne, Switzerland; Inserm U1330, Children’s Oncology Research Unit (CONCERT), Institut Curie, PSL University, Paris, France; Pediatric Hematology-Oncology Unit, Woman-Mother-Child Department, Lausanne University Hospital and University of Lausanne, Lausanne, Switzerland

**Keywords:** Neuroblastoma, ALK activating mutation, xenograft, metastasis, transcriptomic profiling

## Abstract

High-risk neuroblastoma remains a major clinical challenge despite intensive multimodal treatment. Alterations in the anaplastic lymphoma kinase (*ALK*) gene are frequent in NB and correlate with poor outcome. ALK^-F1174L^ and ALK^-R1275Q^ are the most prevalent activating mutations, but their specific effects on tumor behavior remain unclear. Here, we compared the impacts of ALK-wild-type (wt), ALK^-F1174L^ and ALK^-R1275Q^ in several NB xenograft models. In the SK-N-Be2c model, ALK^-F1174L^ mediated numerous transcriptomic alterations without affecting tumor behavior. In GIMEN cells, only ALK^-F1174L^ and ALK^-R1275Q^ conferred tumorigenicity with distinct growth kinetics and transcriptional programs. In the mesenchymal SH-EP model, ALK variants induced distinct phenotypes, with ALK-wt and ALK^-F1174L^ promoting partial transition toward neuroblastic identity. All ALK variants induced lung metastases, but with allele-specific dissemination patterns. Transcriptomic profiling of SH-EP tumors and corresponding metastases revealed partly opposing pathway regulation between ALK^-F1174L^ and ALK^-R1275Q^. Overall, ALK variants differentially modulate NB behavior in a context-dependent manner.

## INTRODUCTION

Neuroblastoma (NB) is a pediatric cancer of the sympathetic nervous system, contributing to 15% of all childhood cancer-related deaths (1). NB exhibits highly variable clinical behaviors, ranging from spontaneous regression to aggressive, treatment-resistant malignancies. Despite intensive multimodal therapies, the overall survival of children with high-risk NB (HR-NB) remains approximately 50% (1). Activating mutations in the anaplastic lymphoma kinase (*ALK*) gene are the most frequent recurrent oncogenic event in NB, identified in both sporadic (6-12%) and familial (50%) NB cases (2–5). *ALK* mutations occur at three major hotspots: R1275 (43%), F1174 (30%), and F1245 (12%). ALK^-F1174L^, restricted to sporadic cases, and ALK^-R1275Q^, the most frequently observed mutation in both familial and sporadic cases, are the most studied mutations (6). ALK^-F1174L^ mediates higher levels of autophosphorylation and kinase activity, increased transforming capacity, and enhanced resistance to first and second-generation ALK inhibitors, compared to ALK^-R1275Q^ and other variants (2, 6–10). Moreover, ALK^-F1174L^ confers higher tumorigenic potential and to cooperate more strongly with MYCN in driving NB development than ALK^-R1275Q^ (11–13). Besides point mutations, *ALK* is altered in NB through copy number gain (17%) or focal genomic amplification (1-2%), the latter almost exclusively co-occurring with *MYCN* amplification (4, 6, 14, 15). *ALK* alterations, through amplification or mutations, are associated with poor outcome in the overall NB cohort and HR-NB (6, 10, 16). Moreover, high expression of wild-type ALK (ALK-wt) in primary NB has been associated with adverse prognostic markers and reduced patient survival (17).

Furthermore, the frequency of *ALK* mutations increases at relapses, arising either *de novo* or through expansion of pre-existing subclones, underscoring *ALK* as an attractive therapeutic target in NB (18, 19). The third-generation inhibitor lorlatinib has shown promising activity across canonical *ALK* mutations and is currently under evaluation in combination with chemotherapy in newly diagnosed HR-NB (14).

Therapeutic response, including sensitivity to ALK inhibitors, is partly influenced by tumor cell plasticity, whereby NB cells transition between noradrenergic (NOR) and mesenchymal (MES) states through epigenetic reprogramming (20–22). Even in the presence of activating mutation, *ALK* is expressed only in the NOR state and repressed in MES cells (22). Whether ALK signaling contributes to the maintenance of the NOR identity or regulates cell-state transitions remain unknown. Additionally, ALK^-F1174L^ and ALK^-R1275Q^ mutations differ in kinase activity and oncogenic potential, but it remains unclear whether they elicit distinct downstream signaling pathways and transcriptional programs. Here, we investigated the impacts of ALK-wt, ALK^-F1174L^, and ALK^-R1275Q^ expression in different cellular contexts using murine xenograft models generated from MYCN-amplified NOR (SK-N-Be2c) and non-MYCN-amplified MES (SHEP and GIMEN) NB cell lines.

## MATERIAL & METHODS

### Cell culture

SK-N-Be2c (RRID:CVCL_0529) (MYCN-amplified), GIMEN (RRID:CVCL_1232) and SH-EP (RRID:CVCL_0524) (MYCN non-amplified) NB cell lines were cultured in Dulbecco’s modified Eagle’s medium (DMEM) (Gibco,Paisley, UK), supplemented with 1% penicillin/streptomycin (Gibco) and 10% heat inactivated Fetal Calf Serum (FCS) (BioWest, South American) and under standard culture conditions in humidified incubator at 37°C with 5% CO2. Cell lines authentication was performed before starting experiments by microsatellite short tandem repeat analysis (Microsynth, Switzerland). Cells were confirmed to be free of mycoplasma contamination using regular PCR testing.

### Plasmid construction and lentiviral infection

The human ALK cDNA *EcoR1-Pme1* fragments, isolated from ALK-wt-, ALK^-F1174L^- and ALK^-R1275Q^-pcDNA3 constructs were introduced into *EcoR1-Pme1* sites of the lentiviral vector pLiVpuro_C located downstream of the EF1-α promoter (23, 24). ALK-wt-, ALK^-F1174L^- and ALK^-R1275Q^-encoding pLiVpuro_C vectors were verified by sequencing of the complete ALK sequence. Lentiviral particles were produced in HEK293T cells by calcium phosphate transfection using empty pLiVpuro_C (Control) or ALK-wt-, ALK^-F1174L^- and ALK^-R1275Q^- pLiVpuro_C vectors together with the packaging plasmids pCMVΔ8 and pMD2. G. Viral infection was performed as previously described (25), except for GIMEN cells which were not supplemented with Polybrene due to its toxicity. Transduced cells were selected using puromycin (Gibco Lifes ® Technologies^TM^) at 5 μg/mL for SK-N-Be2c and 1 μg/mL for SH-EP and GIMEN cells. GIMEN cells were cloned in 96 well plates, and clones were screened by immunocytochemistry (ICC) for ALK expression. Four clones displaying high ALK expression were pooled for each ALK variant. For GIMEN pLiV, 4 clones were randomly selected. Pooled GIMEN cells were subsequently transduced with pLenti-PGK-Venus-Akaluc (neo) (Addgene, plasmid#124701) expressing Venus and luciferase.

### In vivo studies

Animal experiments were carried out in accordance with established guidelines for animal care of the Swiss Animal Protection Ordinance and the Animal Experimentation Ordinance of the Swiss Federal Veterinary Office (FVO). Animal experimentation protocols were approved by the Animal Experimentation Ethics Committee of the Veterinary Service of the Canton of Vaud (Etat de Vaud, Veterinary Service, authorization numbers: VD2995, VD3372 and VD3695). All reasonable efforts were made to reduce suffering.

For orthotopic (ortho) implantations, 5x10^4^ SK-N-Be2c cells were resuspended in 10 μl of PBS and injected in left adrenal gland of athymic Swiss nude mice (Crl:NU(Ico)-Foxn1^nu^, Charles River Laboratory, France) as described (25, 26). Tumor growth was followed by ultrasound every 7 to 14 days and tumor volume (mm^3^) calculated as = 4/3×π×(depth×sagittal×transversal)/6.

For subcutaneous (sc) implantations performed with GIMEN or SH-EP cells, groups of 6 NOD-SCIDγ mice (NSG^TM^, Jackson Laboratory) were injected in the right flank with 2x10^6^ cells in a volume of 200 ul 1:1 mix of DMEM and BD Matrigel^TM^ Basement Membrane Matrix (BD Biosciences, Bedford, MA, USA). Subcutaneous xenografts were also performed athymic Swiss nude mice with 5×10^6^ of SH-EP-ALK-wt, -ALK^-F1174L^, or -ALK^-R1275Q^ cells (6 mice per group). Mice were then monitored twice a week and tumor growth was measured with calipers using the formula: volume (mm^3^) = (length x width^2^)/2.

Mice were euthanized once tumor volumes reached 900 mm^3^ for ortho and 1000 mm^3^ for sc implantations. Tumors and organs were cut into pieces and snap-frozen in liquid nitrogen or fixed in Histofix (ROTI^®^Histofix, Roth AG) and embedded in paraffin.

### Quantification of lung metastases

Intrapulmonary metastases were detected on 3 lung sections (3 µm thick), separated by a depth of 300 µm, by in situ hybridization using the Alu positive probe II (Roche Diagnostics, Cat. No. 05272041001) as described in (25). Whole slides were scanned using the Zeiss Axioscan Z.1 (Zeiss, Oberkochen, Germany) and Alu^+^ cells were quantified using QuPath software (version 0.2.3) at 3X magnification.

### RNA isolation and sequencing

Total RNA from SK-N-Be2c and SH-EP samples were extracted using miRNeasy Mini Kit and RNeasy Mini kit for GIMEN samples (Qiagen, Germany). RNA sequencing was performed at the iGE3 Genomics platform (University of Geneva, https://ige3.genomics.unige.ch/). RNA quantification was performed using a Qubit fluorometer (ThermoFisher Scientific) and RNA integrity assessed using the Agilent 2100 Bioanalyzer system (Agilent Technologies). The Illumina TruSeq Stranded Total RNA library Prep Gold kit was used for the library preparation with 200 ng of total RNA as input (for Experiment 1: SK-N-Be2c and Experiment 2: SH-EP tumors) and the Illumina Stranded Total RNA Prep, Ligation with Ribo-Zero Plus kit with 100 ng of total RNA as input (for Experiment 3: GIMEN and SH-EP lungs and selected SH-EP tumors: T_F1_21, T_F2_21, T_F4_21, T_R2_21, T_R3_21 and T_R5_21). The molarity and quality of the libraries were assessed using Qubit and Tapestation (DNA High sensitivity chip). The libraries were sequenced on an Illumina HiSeq 4000 sequencer (for Exp. 1 & 2) or NovaSeq 6000 (For Exp. 3) sequencer for single-end 100 reads.

### Reads trimming, alignment and gene quantification

To account for the T-overhang due to library preparation, reads were trimmed using Trimmomatic (version 0.39) to remove the first base (HEADCROP:1) for the experiment 2. Reads were aligned using STAR (version 2.5.2b) with a mouse and human combined reference. The reference was built by downloading files from gencode: human GRCh38 genome and mouse GRCm39 genomes in fasta format, with their corresponding (respectively gencode v44 and genocode vM33) transcriptome annotations files in gtf format, all files in the primary assembly version. The main chromosomes were selected, and the chromosomes names were modified in the genomes and annotations to differentiate chromosomes from human and mouse. Alignments from each sample were split into 2 files, one per specie and reads that had at least one alignment for both species were removed. The gene expression was calculated from alignments using RSEM (version 1.3.3). SK-N-Be2c (T_Be) RNAseq samples were analyzed separately, as an independent experiment, compared to SHEP (T_SHEP) and GIMEN (T_GIMEN) samples.

### Gene normalization

The gene normalization was done on R (version 4.2.1), gene counts were rounded (base package, function round) then the lowly filtered genes were removed by keeping only genes with at least 10 raw counts in at least 4 samples. The filtered counts were normalized using median of ratios method (DESeq2 package version 1.38.3, function estimateSizeFactors). Finally, normalized counts were log2-transformed using the log2 function (base package) with a pseudocount of 1.

### Mutated allele fraction determination in Be2C

Using the RNAseq reads aligned to ALK in the expected mutation sites, the mutated allele fraction (MAF) was calculated for each sample and each position by dividing the number of reads with mutated allele by the number of reads aligned.

### Differential gene expression analysis

In R (version 4.2.1), the expected gene counts were rounded (base package, function round) and the lowly filtered genes were removed by keeping only genes with at least 10 raw counts in 4 samples. Differentially expressed genes were calculated using DESeq2 package (version 1.38.3, DESeq function). The cutoffs for log2 fold-change and adjusted p-values to call differentially expressed genes were set to 1 and 0.05, respectively.

### Enrichment analysis

Over representation analysis (ORA) was performed on differentially expressed genes using enrichGO function of clusterProfiler package (version 4.6.2) for GO database (version 2022-07-01) and enrichPathway function of ReactomePA package (version 1.42.0) for Reactome database. For each comparison the analysis was performed separately on the up-regulated, down-regulated, and both up- and down-regulated differentially expressed genes. A cutoff of 0.05 was used for adjusted p-value to define enriched pathways. For analyses using Reactome, the gene identifiers were converted to ENTREZID using gconvert function (gProfiler2 package version 0.2.3, with the mthreshold=1 option to get only one identifier when there are multiple).

### Signature scoring

Rank-based pathway activity score was calculated on log2-transformed normalized counts. The scoring was implemented in Perl (version 5.34.0) as previously described (27). The R package singscore was used to obtain sample-wise, cohort-independent scores (28).

### Statistics and reproducibility

All statistical analyses (except for transcriptomic analyses) were performed using GraphPadPrism 8.3.0 (GraphPad Software Inc., San Diego, CA, USA). D’Agostino-Pearson normality test was performed for each data set. Data were analyzed with unpaired two-tailed parametric t-test or non-parametric Mann−Whitney test depending on distribution to compare two different conditions or as indicated in the Figure legends. RNAseq analyses were performed using 4 tumors or lung tissue fragments per group, except for SH-EP ALK^-F1174L^ and ALK^-R1275Q^ tumors for which 2 additional tumors were added in a second sequencing analysis together with 2 tumors from the first analysis. The *in vivo* studies using SH-EP cells were conducted in NSG mice and once in nude mice. RT-qPCR analyses were performed on 6 tumors and 3 lung tissues per mice group. IHC was performed on all available tumor and lungs tissues (n=6). Additional methods are described in the Supplementary Methods, including the list of primers and antibodies sources and dilutions.

## RESULTS

### ALK variant overexpression does not alter SK-N-Be2c cell tumorigenicity or phenotype despite distinct transcriptomic alterations

The SK-N-Be2c cell line was initially selected among NOR cell lines for ALK overexpression and *in vivo* studies because it weakly expresses the endogenous ALK gene (wt genotype) (Suppl. Fig. 1A, 1B). SK-N-Be2c control (pLIV) and ALK-wt, ALK^-F1174L^ and ALK^-R1275Q^ - overexpressing cells were orthotopically injected into athymic Swiss nude mice. All tumor groups had similar growth kinetics and histological features, consisting of stroma-poor NB phenotype with undifferentiated or poorly differentiated neuroblasts (Fig. 1A, Suppl. Fig. 1C). Endogenous *ALK* expression was detected in SK-N-Be2c-pLIV control tumors, with both intra- and inter-tumoral heterogeneity. ALK expression was increased in ALK-wt, ALK^-F1174L^ and ALK^-R1275Q^ tumors relative to controls, with ALK^-R1275Q^ showing the lowest levels among ALK transduced tumor groups (Suppl. Fig. 1C, 1D).

**Figure 1.**
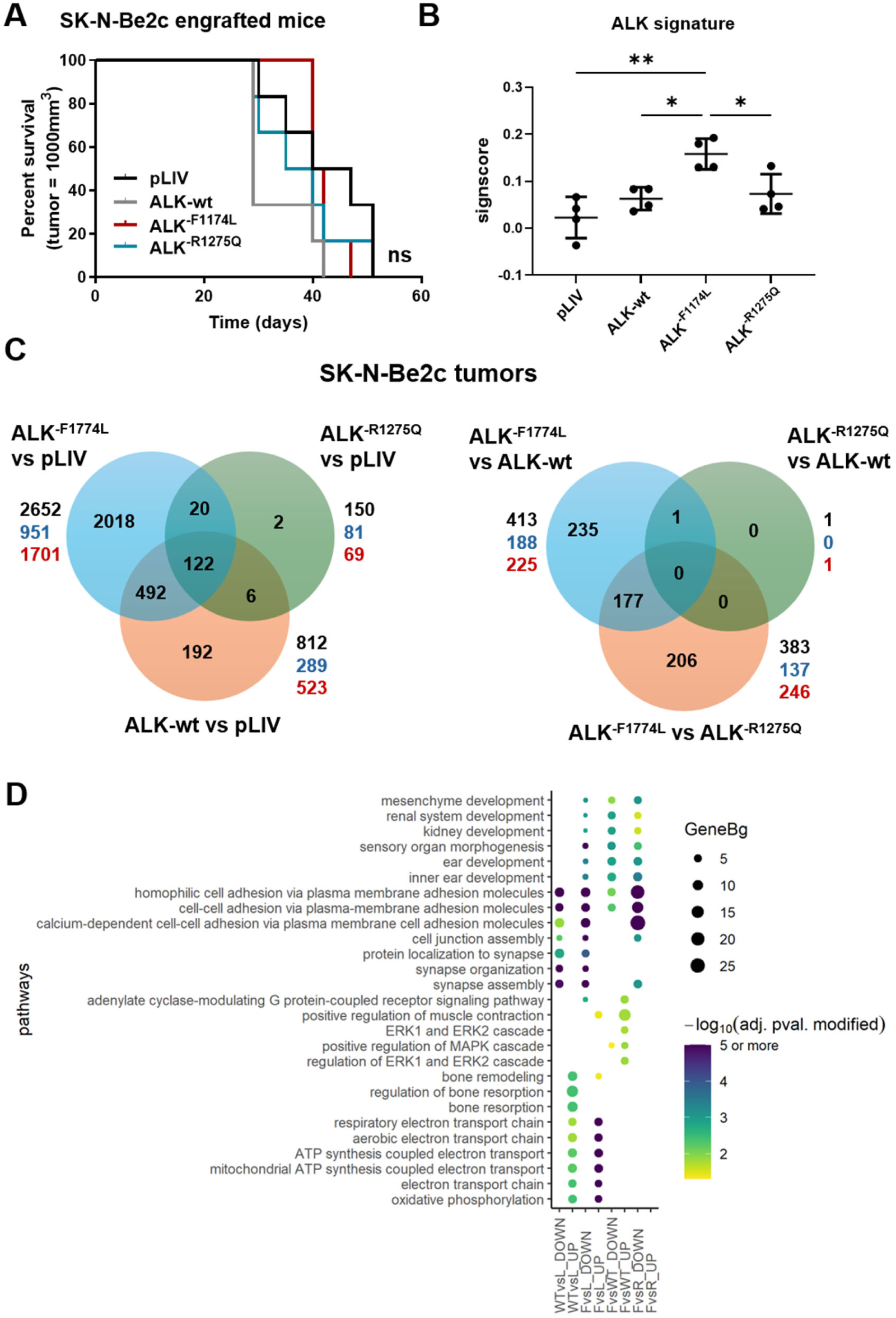
Impact of ALK-wt, ALK^-F1174L^ or ALK^-R1275Q^ overexpression in the SK-N-Be2c orthotopic model. **(A)** Kaplan-Meier survival curves of athymic Swiss nude mice implanted orthotopically with SK-N-Be2c Control (pLIV) and ALK-expressing cells (n=6 mice/group). Log rank test curve comparisons, ns: not significant. **(B)** ALK signature scores calculated on protein coding genes expressed in SK-N-Be2c tumors are plotted as mean ±SD. One-way Anova test: * *p*<0.05, \*\**p*=0.001. **(C)** Venn diagram showing the common and specific DE genes (adj p-value <0.05 et absolute log2FC ≥1) in SK-N-Be2c ALK transduced-derived tumors vs pLIV control tumors (upper panel) or between each ALK-overexpressing tumor (lower panel). Total number of DE (black), upregulated (blue) and downregulated (red) genes are also indicated for each comparison. **(D)** Dot plot of the top 5 (by adjusted p-values) enriched GO BP pathways among upregulated (UP) and downregulated (DOWN) gene lists in each indicated comparison of SK-N-Be2c tumors. Dot size represents the enrichment factor measured as Gene ratio/background ratio (GeneBg), and the color indicates the significance level (adjusted p-value, −log10 scale). L = pLIV, WT = ALK-wt, F = ALK^-F1174L^, R = ALK^-R1275Q^.

Transcriptomic analyses of the orthotopic tumors confirmed a predominant expression of the mutant alleles in ALK^-F1174L^ and ALK^-R1275Q^ tumors, with mean mutated allele fractions of 92% and 76%, respectively (Suppl. Fig. 1E). To further validate ALK signaling activity within the tumors, we analyzed the 77-ALK gene signature (29). ALK activity score was significantly increased in ALK^-F1174L^ tumors, while ALK^-R1275Q^ and ALK-wt tumors showed a non-significant upward trend vs pLIV tumors (Fig. 1B).

Compared with pLIV tumors, ALK^-F1174L^ tumors showed more transcriptional alterations (2652 differentially expressed genes, DEGs), followed by ALK-wt tumors (812 DEGs), with most genes being downregulated in both comparisons, while ALK^-R1275Q^ tumors displayed only 150 DEGs. *ALK* was the only DEG identified when comparing ALK-wt versus (vs) ALK^-R1275Q^ tumors. In contrast, ALK^-F1174L^ vs ALK^-R1275Q^ and ALK^-F1174L^ vs ALK-wt comparisons revealed similar numbers of DEGs (383 and 413, respectively), although fewer than half were shared between the two comparisons, indicating distinct transcriptomic programs associated with each ALK variant. (Fig. 1C, Suppl. Data 1).

Main pathways enriched in Gene Ontology (GO) biological process (BP) and REACTOME databases for the downregulated DEGs in ALK^-F1174L^ and ALK-wt tumors vs pLIV tumors, were involved in cell-cell adhesion, synapse assembly and organization, which include several protocadherins and cadherins genes, and neuronal development (Fig. 1D, Suppl. Fig.1F, Suppl. Data 2 and 3). In contrast, pathways enriched in upregulated DEGs in ALK^-F1174L^ and ALK-wt tumors vs pLIV tumors were predominantly associated with oxidative phosphorylation and nucleotide biosynthesis. Moreover, pathways associated with tissue and bone resorption were specifically enriched in upregulated DEGs in ALK-wt vs pLIV tumors (Fig. 1D, Suppl. Data 2). Additionally, developmental pathways were enriched in the downregulated DEGs in ALK^-F1174L^ tumors vs all other groups, whereas pathways associated with the regulation of the MAPK (ERK1/2) signaling cascade were enriched in ALK^-F1174L^ vs ALK-wt upregulated DEGs (Fig. 1D, Suppl. Data 2). Altogether, these results highlight the strong impact mediated by the ALK^-F1174L^ mutation in the SK-N-Be2c genetic background and suggest a possible distinct impact of ALK-wt and ALK^-F1174L^ variants on tumor transcriptomic profiles. Nevertheless, endogenous *ALK*-wt expression detected in control tumors, together with its contribution to ∼25% of *ALK* transcripts in ALK^-R1275Q^ tumors, limited the ability to clearly characterize the specific impact of ALK-wt and ALK^-R1275Q^ isoforms in this model.

### ALK^-F1174L^ and ALK^-R1275Q^ mutations confer tumorigenicity to GIMEN cells and drive distinct transcriptomic programs

To further investigate the distinct effect of ALK variants *in vivo*, we selected the weakly tumorigenic and MES GIMEN and SH-EP NB cell lines, lacking ALK expression *in vitro* (Suppl. Fig. 1A). GIMEN-pLIV and ALK-wt-, ALK^-F1174L^-, and ALK^-R1275Q^-expressing cells were subcutaneously implanted into NOD-SCID-*γ* (NSG) mice. None of the mice implanted with GIMEN-pLIV or -ALK-wt developed tumors within the 195-day experiment. In contrast, all mice implanted with GIMEN-ALK^-F1174L^ (6/6) or -ALK^-R1275Q^ (5/5) developed sc tumors, with a faster growth observed in the ALK^-R1275Q^ group (Fig. 2A). Histologically, tumors displayed a heterogeneous composition, with regions presenting elongated, MES-like cells enriched in extracellular matrix (ECM) components, along with regions composed of small round neuroblast-like cells. MES-like cells were more abundant in ALK^-F1174L^ tumors, whereas ALK-^R1275Q^ tumors predominantly displayed neuroblast-like cells (Fig. 2B). ALK was strongly expressed in all transduced GIMEN cells and derived tumors (Suppl. Fig. 2A, Fig. 2B). The high expression of ALK-wt alone was insufficient to drive tumor formation, potentially due to the lack of ligand-mediated activation or insufficient expression level for autoactivation (17).

**Figure 2.**
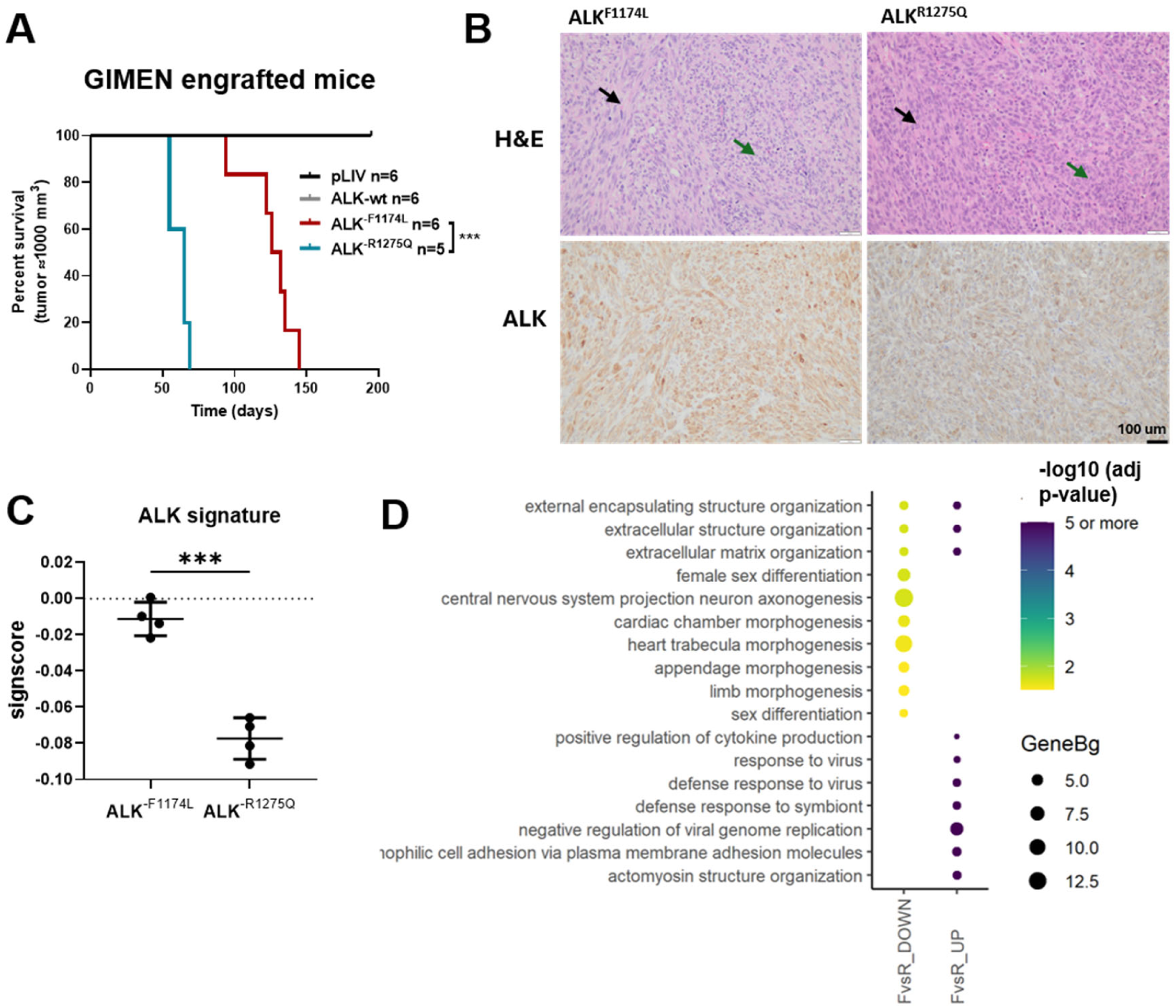
ALK^-F1174L^ and ALK^-R1275Q^ mutations confer tumorigenicity to GIMEN cells with distinct growth kinetics and transcriptomic features. **(A)** Kaplan-Meier survival curve of mice injected with GIMEN cell variants, showing cumulative survival probability throughout the follow-up period. Median: 65 days for ALK^-R1275Q^ group, 129 days for ALK^-F1174L^ group. Log-rank Mantel-Cox test ***p<0.001. *n* = 6 for pLIV, WT, and ALK^-F1174L^; *n* = 5 for ALK^-R1275Q^, as one mouse developed a spontaneous thymic tumor and was excluded from the analysis. **(B)** Representative images of H&E and ALK staining in GIMEN-derived tumors. Black arrows indicate elongated, MES-like cells, while green arrows point to small, round neuroblast-like cells. **(C)** ALK signature scores calculated on protein coding genes expressed in GIMEN tumors are plotted as mean ±SD. Unpaired t-test: * *p*<0.05, \*\*\**p*=0.0001. **(D)** Dot plot showing the top 10 enriched GO BP sorted by adjusted p-values among upregulated and downregulated genes in GIMEN tumors. Dot size represents the enrichment factor measured as Gene ratio/background ratio (GeneBg), and the color indicates the significance level (adjusted p-value, −log10 scale). F = ALK^-F1174L^, R = ALK-R1275Q.

Furthermore, transcriptomic analysis identified 1964 DEGs between ALK^-F1174L^ and ALK^-R1275Q^ tumors, indicating an activation of distinct transcriptomic programs (Suppl. Fig. 2B, Suppl. Data 4). As in the SK-N-Be2c model, ALK^-F1174L^ tumors exhibited higher ALK signature scores than ALK^-R1275Q^ tumors, confirming a stronger signaling activity despite slower growth (Fig. 2C). Enrichment was limited among downregulated DEGs, including developmental processes, such as morphogenesis and neuron axonogenesis (Fig. 2D, Suppl. Data 5 and 6). In contrast, numerous pathways were enriched among upregulated DEGs in ALK^-F1174L^ vs ALK^-R1275Q^ tumors, including cytokine production, antiviral response, and interferon signaling, suggesting a tumor cell-intrinsic activation of stress response and inflammatory signaling, as well as extracellular matrix organization, cell adhesion, ECM interaction (Fig. 2D, Suppl. Fig.2C, Suppl. Data 5 and 6). These pathways enrichment are consistent with the MES-like phenotype observed in ALK^-F1174L^ tumors and might explain their slower growth.

### ALK variants differentially alter tumor phenotype and metastasis in the SH-EP model

All NSG mice subcutaneously engrafted with SH-EP-pLIV, -ALK-wt, -ALK^-F1174L^-, or -ALK^-R1275Q^ cells developed tumors (6/6 per group), although growth dynamics varied according to ALK variants. ALK^-F1174L^ markedly accelerated tumor growth, reducing median survival to 47 days compared with 83 days for controls and 90 days for ALK^-R1275Q^ tumors, whereas ALK-wt had a moderate effect (71 days) (Fig. 3A). All SH-EP-ALK cells and their derived tumors expressed ALK, while controls lacked endogenous *ALK* expression (Suppl. Fig. 3A, 3B). Histological analysis showed that SH-EP-pLIV and ALK^-R1275Q^ tumors retained a MES phenotype, characterized by fusiform cells with large cytoplasm and a collagen-rich stroma (Fig. 3B). In contrast, ALK-wt and ALK^-F1174L^ tumors were predominantly composed of small round blue cells, resembling poorly differentiated NB, and suggesting a shift toward an undifferentiated, neuroblastic phenotype. In addition, a subset of cells in ALK-wt tumors displayed differentiation toward ganglion-like cells (Fig. 3B). This phenotypic change was further supported by a focal loss of the MES marker CD44 in distinct tumor regions (Fig. 3C). Interestingly, 2/6 ALK-wt tumors contained regions with PHOX2B^+^ cells (Fig. 3C). Similar phenotypic changes and CD44 expression were also observed in athymic Swiss nude mice (Suppl. Fig. 3C). These findings suggest that ALK-wt and ALK^-F1174L^ reduce the MES phenotype and promote a transition toward a neuroblastic cell identity.

**Figure 3.**
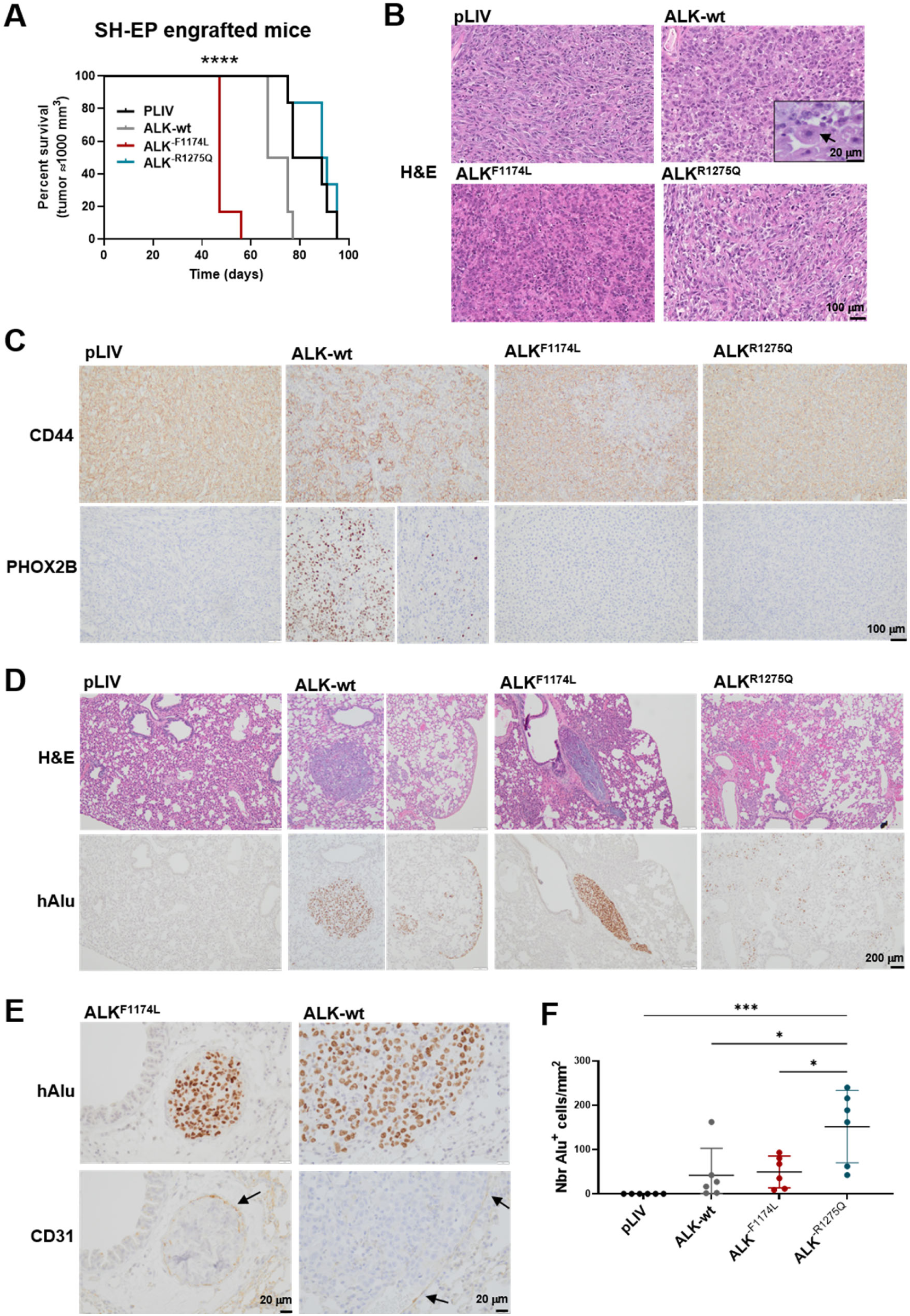
ALK variants induced distinct tumor phenotypes and metastatic patterns in the SH-EP model. **(A)** Kaplan-Meier survival curve of mice injected with SH-EP cell variants (n= 6 mice/group), showing cumulative survival probability throughout the follow-up period. Mice were sacrificed once tumor reached 1000 mm^3^. Log-rank Mantel-Cox tests: all groups: ****p<0.0001, FvsR: \*\*\**p*=0.0005, FvsWT: \*\*\**p*=0.0005, FvspLIV: \*\*\**p*=0.0005, WTvsR: \*\**p*=0.0029, WTvspLIV: \**p*=0.0102, RvspLIV: *p*=0.4179. **(B)** Representative images of H&E staining in SH-EP derived tumors illustrating overall tissue architecture. Scale bars: 100 and 20 μm **(C)** Representative images of CD44 and PHOX2B IHC staining showing changes in mesenchymal and noradrenergic marker expression, respectively, depending on ALK variants expression. Scale bar: 100 μm. **(D)** Representative images of lung sections showing heterogeneous dissemination patterns of SH-EP tumor cells detected using human Alu in situ hybridization (ISH) and H&E staining. Scale bar: 200 μm. **(E)** Tumor embolus detected by hAlu ISH and anti-CD31 IHC (indicated by arrow) in lung from a mouse injected with SH-EP ALK^-F1174L^ or ALK-wt cells. Scale bar: 20 μm. **(F)** Quantification of the number of disseminated h-Alu+ cells in the lungs was performed using the QuPath software. Dot plot showing the total numbers of h-Alu+ cells detected on 3 lung slides separated by 300 μm per mm^2^ of lung surface analyzed for each mouse, with mean ±SD (n=6 mice/group). Ordinary one-way ANOVA, *p <0.02, ***p =0.0005.

We next investigated the occurrence of lung metastasis in mice engrafted with SH-EP cells. All mice implanted with SH-EP-ALK expressing cells exhibited disseminated tumor cells (DTCs), whereas none were detected in control mice (Fig. 3D). Moreover, distinct metastatic patterns were observed between ALK variants. Specifically, ALK^-R1275Q^ tumors generated numerous isolated or small DTC clusters within the lung parenchyma, whereas ALK^-F1174L^ tumors mainly formed limited number of large intravascular tumor emboli (ITE) (>1000 mm^2^) with few extravascular DTCs (Fig. 3D, 3E). Notably, mice implanted with SH-EP-ALK-wt cells showed mixed dissemination patterns, including few isolated or small DTC clusters, rare ITE, and established metastases in the peripheral regions of the lung parenchyma (Fig. 3D and 3E). Quantification of Alu-positive cells confirmed increased dissemination in the ALK^-R1275Q^ group (Fig. 3F). Together, these distinct metastatic behaviors, along with the heterogeneous phenotypic changes observed in the tumors, suggest that ALK-wt, ALK^-F1174L^ and ALK^-R1275Q^ drive different signaling outputs.

### ALK variants drive distinct transcriptomic alterations in SH-EP-derived tumors

RNA-sequencing analyses were performed on all groups of SH-EP-derived tumors and on lung tissues isolated from the ALK^-F1174L^ and ALK^-R1275Q^ groups. Unsupervised hierarchical clustering analysis revealed that ALK-wt, ALK^-F1174L^ and pLIV samples were more closely related to each other than to ALK^-R1275Q^ tumors and lung tissues (except for T_SHEP_R6), suggesting that ALK^-R1275Q^ samples exhibit a more distinct gene expression pattern (Suppl. Fig. 4A).

DE analysis across SH-EP tumor groups revealed extensive transcriptional differences, with the highest number of DEGs identified between ALK^-R1275Q^ and either ALK^-F1174L^ or ALK-wt tumors (4466 and 3857 DEGs, respectively), while ALK^-F1174L^ vs ALK-wt tumors were more similar (1433 DEGs) (Suppl. Fig. 4B, Suppl. Data 4). Consistently, ALK-wt and ALK^-F1174L^ tumors displayed numerous DEGs relative to control tumors (n=2963 and n=3327, respectively) and shared the largest overlap of DEGs (n=1713), with only 1% showing opposite fold changes (FC) (Suppl. Fig. 4B, and 4C, Suppl. Data 4). In contrast, ALK^-R1275Q^ tumors presented fewer transcriptomic changes relative to pLIV tumors (2272 DEGs), a smaller overlap with either ALK^-F1174L^ (n=947) or ALK-wt (n=847), and a higher proportion of genes showing opposite FC (8 to 10%, respectively), further supporting the distinct transcriptional program associated with this variant (Suppl. Fig. 4B and 4C, Suppl. Data 4).

**Figure 4.**
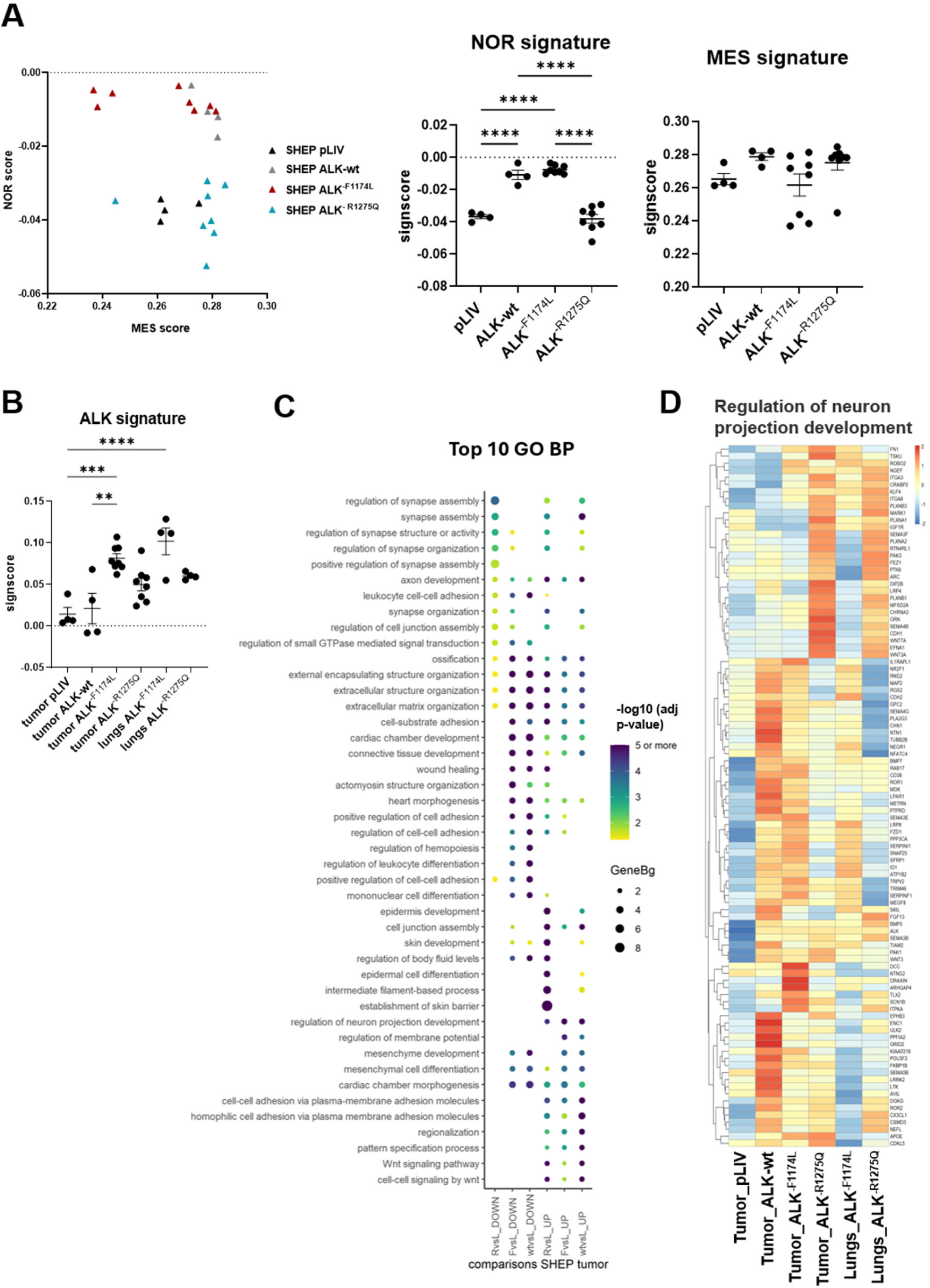
**(A)** Graphs showing the MES and NOR signature scores of SH-EP derived tumors plotted together or as individual values with mean ±SD **(B)** ALK signature scores in SH-EP tumor and lung samples are plotted with mean ±SD. (A-B) Signature scores were calculated on protein coding genes. One-way Anova test: * *p*<0.05, \*\*\**p*=0.0002, ****p<0.0001. **(C)** Dot plot showing the top 5 or 10 enriched GO BP pathways sorted by adjusted p-values among upregulated and downregulated genes in the diverse comparisons of SH-EP tumors. Dot size represents the enrichment factor measured as Gene ratio/background ratio (GeneBg), and the color indicates the significance level (adjusted p-value, −log10 scale). L = pLIV, WT = ALK-wt, F = ALK^-F1174L^, R = ALK^-R1275Q^, UP = upregulated, DOWN = downregulated. **(D)** Heatmap of the Regulation of neuron projection development GO BP pathway representing only the DEGs of ALK expressing tumors vs pLIV tumors. Z-score scaled by row is shown for each sample group.

Moreover, analysis of murine DEGs revealed that ALK^-R1275Q^ tumors displayed the most divergent stromal transcriptomic landscape, differing from ALK^-F1174L^ and ALK-wt tumors with 1186 and 852 DEGs, respectively, while ALK-wt and ALK^-F1174L^ tumors remained highly similar (15 DEGs) (Suppl. Data 7). Together, these findings suggest that the molecular alterations associated with ALK-wt, ALK^-F1174L,^ and ALK^-R1275Q^ signaling in SH-EP tumors are only partially overlapping, with ALK^-R1275Q^ driving a more distinct transcriptional program in both tumor and stromal cells.

To validate the changes in tumor phenotype observed by IHC, and assess a potential shift in cell state, the MES and NOR signature scores were analyzed (20) (adrenergic (ADR) is referred to as NOR in this manuscript). SH-EP-pLIV and -ALK^-R1275Q^ tumors clustered together and showed lower NOR scores relative to SH-EP-ALK-wt and -ALK-F^-F1174L^ tumors. Notably, three SH-EP-ALK^-F1174L^ tumors also showed reduced MES scores, suggesting a stronger shift from MES toward NOR state (Fig.4A). As observed in SK-N-Be2c and GIMEN models, ALK^-F1174L^ tumors showed the highest ALK signature score (Fig. 4B).

Pathways of cell-cell adhesion via plasma membrane adhesion molecules, cell junction assembly, Wnt signaling, neuronal and axon development were enriched in the upregulated genes in all ALK-expressing tumors relative to pLIV controls (Fig. 4C and Suppl. Data 5 and 6). However, pathways associated with mesenchymal development and differentiation were specifically enriched in the DEGs of ALK^-F1174L^ and ALK-wt relative to pLIV tumors, consistent with the altered MES cell phenotype observed in these tumor groups (Fig. 4C, 3B and 4A). Several pathways such as wound healing and ECM proteoglycan displayed opposite enrichment patterns, being enriched in the upregulated DEGs in ALK^-R1275Q^ and in the downregulated DEGs ALK^-F1174L^ and ALK-wt relative to pLIV tumors. Other pathways associated with extracellular matrix, cell substrate adhesion, collagen formation, were enriched in upregulated genes in ALK^-R1275Q^ vs pLIV tumors but were globally dysregulated in ALK^-F1174L^ and ALK-wt tumors. Furthermore, pathways related to skin development were uniquely enriched among the upregulated DEGs in ALK^-R1275Q^ tumors vs pLIV, while pathways linked to synapse organization and assembly were globally dysregulated in this group (Fig. 4C and Suppl. Data 5 and 6). Heatmaps of DEGs within enriched pathways further illustrate that each ALK variant dysregulated distinct gene sets, with particularly pronounced divergence observed in ALK^-R1275Q^ tumors (Fig. 4D, Suppl. Fig. 4E).

To better understand the effects of ALK variants on ALK signature score, DEGs associated with the ALK signature were examined. Heatmap representing DEGs across tumors revealed several genes regulated in the opposite direction compared to the consensus signature (Suppl. Fig. 5A). Both ALK-wt and ALK^-F1174L^ induced a higher level of ETV transcription factors relative to ALK^-R1275Q^, including *ETV1* and *ETV5,* as well as *ETV4* which is not part of the canonical ALK signature (Suppl. Fig. 5B). Similarly, the MAPK pathway feedback regulators *SPRY2* and *SPRY4* were more strongly induced by ALK-wt and ALK^-F1174L^, whereas *DUSP6* was upregulated by all ALK variants (Suppl. Fig. 5B). The mRNA expression levels of selected genes, frequently identified across enriched pathways, were also measured by RTqPCR in SH-EP tumors. *COL4A2, ITGAV* and *SEMA5A* were downregulated, while *SEMA3E* was upregulated in ALK-wt and ALK^-F1174L^ tumors relative to controls. In contrast, the expression of these genes remained unchanged in ALK^-R1275Q^ tumors. *SEMA3B* was upregulated by the three ALK variants, whereas *PLXNA2* upregulation was specific to ALK^-R1275Q^ tumors (Suppl. Fig. 5B). Altogether, these findings indicate that each ALK variant mediates partially distinct transcriptomic alterations in the SH-EP background. ALK-wt exhibits greater similarity to ALK^-F1174L^, whereas ALK^-R1275Q^ displayed the most divergent transcriptional profile. The distinct pathway enrichment patterns and differential expression of specific gene sets underscore the variant-specific regulatory effects of ALK signaling.

### ALK^-F1174L^ and ALK^-R1275Q^ induce distinct metastatic transcriptional programs

To investigate transcriptomic remodeling associated with the distinct dissemination patterns mediated by ALK^-F1174L^ and ALK^-R1275Q^ mutations, lung lesions were compared with primary tumors in each mouse group. This analysis identified 635 DEGs in the ALK^-F1174L^ group and 1501 in the ALK^-R1275Q^ group, with only 98 overlapping genes, of which 22% displayed opposite FC (Fig. 5A). Pathways related to ECM/extracellular structure organization, collagen metabolic process, and axon development represented the top enriched pathways (lowest adjusted p-value) among downregulated DEGs in lung tissue from the ALK^-R1275Q^ group, whereas they showed weaker enrichment among upregulated DEGs in the ALK^-F1174L^ group (Fig. 5B and Suppl. Data 5 and 6). In contrast, pathways related to cardiac development, cell growth, cell and tissue migration, and morphogenesis were top-enriched pathways among upregulated DEGs in ALK^-F1174L^ metastases but showed lower enrichment significance among downregulated DEGs in ALK^-R1275Q^ disseminated cells (Fig. 5B and Suppl. Data 5). These results are consistent with the cohesive architecture of tumor emboli and reflect activation of embryonic morphogenetic programs that support collective migration behavior. In ALK^-R1275Q^ lung metastases, metabolic pathways linked to mitochondrial respiration and the electron transport chain were enriched (Fig. 5B and Suppl. Data 5 and 6). These observations are consistent with ALK-R1275Q leading to disseminated cells, which may have adopted a metabolically active state with reduced ECM dependency, favoring their survival in the circulation and their adaptation to early metastatic niches. Together, these findings indicate that ALK^-F1174L^ and ALK^-R1275Q^ mutations generate distinct, partly opposing transcriptional programs in lung metastases relative to their respective primary tumors, consistent with their divergent metastatic patterns.

**Figure 5.**
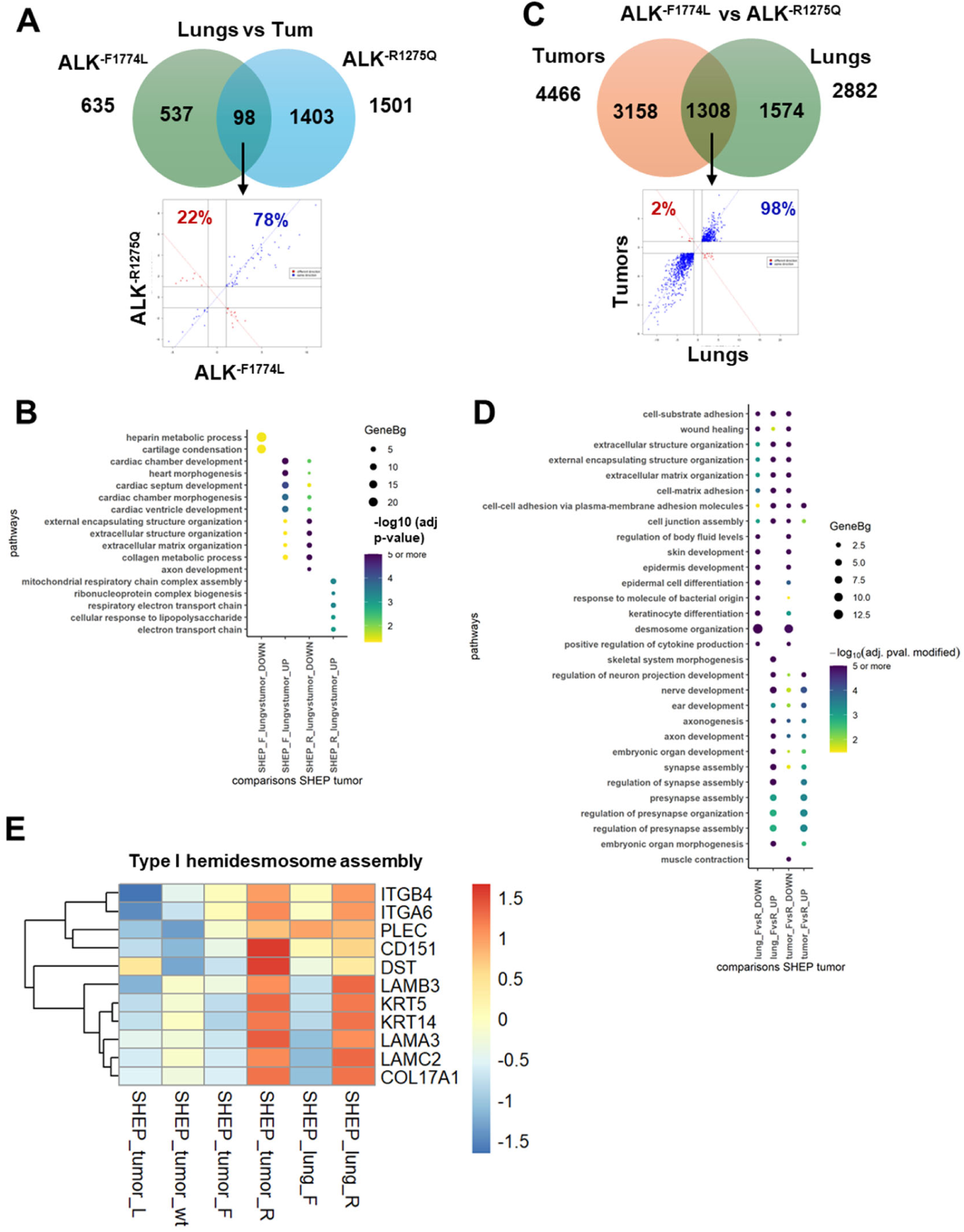
**(A)** Venn diagram showing the common and specific DEGs obtained in Lungs vs Tumors comparisons for the ALK^-F1174L^ or ALK^-R1275Q^ groups and graphs showing the genes deregulated in common for the indicated comparisons plotted according to their log2(FC) and labelled in blue if deregulated in the same direction or in red if deregulated in opposite direction. **(B)** Dot plot showing the top 5 enriched GO BP pathways sorted by adjusted p-values among upregulated and downregulated genes in Lungs vs Tumors comparisons for ALK^-F1174L^ or ALK^-R1275Q^ groups. Dot size represents the enrichment factor measured as Gene ratio/background ratio (GeneBg), and the color indicates the significance level (adjusted p-value, −log10 scale). F = ALK^-F1174L^, R = ALK^-R1275Q^, UP = upregulated, DOWN = downregulated. **(C)** Venn diagram showing the common and specific DEGs between SH-EP cells expressing ALK^-F1174L^ vs ALK^-R1275Q^ in lungs or in tumors and graph of common genes as in (A). **(D)** Dot plot showing the top 10 enriched GO BP pathways sorted by adjusted p-values among upregulated and downregulated genes in ALK^-F1174L^ vs ALK^-R1275Q^ comparisons in lungs or in tumors, as in (B). **(E)** Heatmap of Type I hemidesmosome assembly REACTOME pathway showing the DEGs all group comparisons described in this study. Z-score scaled by row are shown for each sample group. L = pLIV, WT = ALK-wt, F = ALK^-F1174L^, R = ALK^-R1275Q^.

### The differential transcriptional program mediated by ALK^-F1174L^ and ALK^-R1275Q^ mutations is partly conserved in tumors and lung metastases

To determine whether the differences observed in primary tumors were conserved in the disseminated cells, the transcriptomic profiles of ALK^-F1174L^ and ALK^-R1275Q^ lung metastases were compared. Numerous DEGs were detected between ALK^-F1174L^ and ALK^-R1275Q^ groups in lung metastases (n=2882), as observed in primary tumors (n=4466), with 1308 shared DEGs, of which 98% showed the same direction of change (Fig. 5C). These results confirm that the two activating ALK mutations drive distinct transcriptional programs that are partially conserved in metastases.

Furthermore, pathways related to neuronal development and synaptic structure and function were the most significantly enriched pathways in upregulated DEGs in ALK^-F1174L^ vs ALK^-R1275Q^ tumors (Fig. 5D, Suppl. Data 5). Especially, the same pathways were enriched in upregulated DEGs in ALK^-F1174L^ vs ALK^-R1275Q^ lung metastases, indicating that ALK^-F1174L^-mediated transcriptional changes related to neuronal differentiation in primary tumors persist at metastatic site. In contrast, pathways associated with cell adhesion and tissue remodeling were predominantly enriched in downregulated DEGs in SH-EP-ALK^-F1174L^ vs ALK^-R1275Q^ tumors (Fig. 5D, Suppl. Fig. 6A, Suppl. Data 5 and 6). Moreover, several adhesion- and ECM-related pathways (epidermis development, skin development, desmosome organization) and immune-related processes (leukocyte adhesion, cytokine production, regulation of immune cells), remained enriched in downregulated DEGs in ALK^-F1174L^ lung metastases (Fig. 5D). The genes of type I hemidesmosome assembly pathway showed strong upregulation in ALK^-R1275Q^ tumors and lung lesions (Fig. 5E). In contrast, pathways related to extracellular matrix and cell adhesion were enriched in both upregulated and downregulated DEGs in ALK^-F1174L^ vs ALK^-R1275Q^ metastases, with stronger statistical significance in the upregulated DEGs (Fig. 5D, Suppl. Data 5 and 6). The differentially expressed genes of the Reactome non-integrin membrane-ECM pathway further illustrate the difference across all group comparisons (Suppl. Fig. 6B). These data confirm the distinct transcriptomic program mediated by each activating mutation.

To decipher the impact of ALK^-R1275Q^, we focused on the pathways related to desmosomes and hemidesmosomes, which showed the highest gene enrichment ratios in SHEP ALK^-R1275Q^ tumors and lung metastases compared with all other groups. We identified *KTR5*, *KRT14*, *COL17A*, *LAMC2*, *DSP,* and *DSG2*, ranking among the top upregulated DEGs in ALK^-R1275Q^ tumors and lungs, with *KRT14* and *KRT5* displaying the highest log2(FC) (Suppl. Data 4). *KRT14*, the major intermediate filament in hemidesmosomes pairing with *KRT5*, has been described as highly expressed in micrometastases but reduced in macrometastases in breast cancer, where a KRT14 signature of 239 genes has been established (30, 31). Moreover, *KRT14* expression was shown to be required for distant metastasis formation and correlated with a gene network involved in metastatic regulation (30). We confirmed the upregulation of *KRT14, KRT5*, *COL17A,* and *DSP* mRNA in tumors and lung tissues of ALR^-R1275Q^ groups, whereas their expression was almost undetectable in other groups (Suppl. Fig. 7A and 7B). Moreover, KRT14 protein was uniformly expressed in ALK^-R1275Q^ tumors, in most tumor cells, as illustrated by the co-staining with human Alu ISH (Fig. 6A). In contrast, only rare KRT14^+^ cells were detected in ALK-wt tumors, and almost none in pLIV and ALK^-F1174L^ tumors (Fig. 6A). Consistently, KRT14^+^ cells were only detected in the lungs of the ALK^-R1275Q^ group (Fig. 6A). Similarly, KRT5 was strongly expressed in ALK^-R1275Q^ tumors, compared to ALK-wt and ALK^-F1174L^, while it was not detected in pLIV tumors (Fig. 6B). Finally, the KRT14 signature score (30) was significantly higher in ALK^-R1275Q^ tumors compared to other tumor groups, as well as in lung lesions of the ALK^-R1275Q^ group compared to ALK^-F1174L^ and control tumors (Fig. 6C). Altogether, these data reveal that the distinct signaling and transcriptional programs mediated by each ALK activating mutation in the SH-EP model are partly conserved in primary tumors and metastases, with ALK^-R1275Q^ upregulating hemidesmosome and *KRT14*-associated genes.

**Figure 6.**
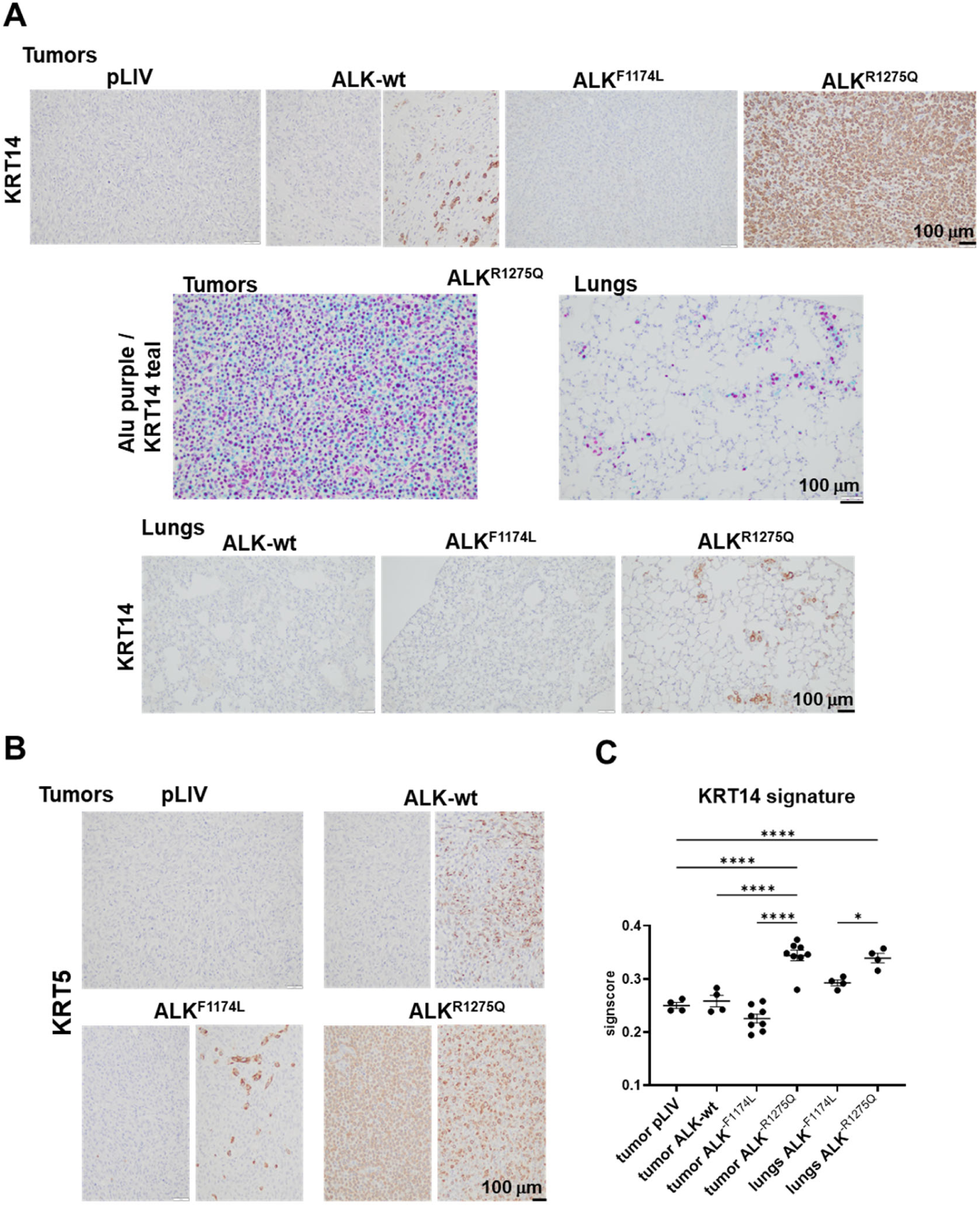
SH-EP-ALK^-R1275Q^ tumors and disseminated cells in the lungs displayed elevated KRT14 and KRT5 expression. **(A)** Representative images of KRT14 expression analyzed by IHC in SH-EP tumors and lungs and of KRT14 or ALU/KRT14 co-staining for ALK^-R1275Q^ group (Alu in purple and KRT14 in teal). Scale bar = 100 μm. **(B)** Representative images of KRT5 expression analyzed by IHC in SH-EP tumors. Scale bar = 100 μm. **(C)** KRT14 signature scores calculated on protein coding genes expressed in SHEP tumors and lungs, and plotted with mean ±SD. One-way Anova test: \**p*<0.05, \*\*\*\**p*<0.0001.

## DISCUSSION

We found that ALK signaling in NB is dependent on both the specific variant and the cellular context. In the SK-N-Be2c model, overexpression of ALK-wt or activating ALK mutations did not enhance tumor aggressiveness, consistent with the highly tumorigenic nature of this cell line (25). Baseline endogenous ALK-wt expression in control and ALK^-R1275Q^ tumors further limited the ability to clearly discriminate variant-specific effects, although ALK^-F1174L^ induced the strongest transcriptomic alterations.

In MES models, we found that ALK overexpression elicited distinct oncogenic effects depending on ALK variants and cellular context. In both GIMEN and SH-EP models, tumors retained an overall MES phenotype. This is consistent with previous data showing that tumors derived from SH-SY-5Y cells expressing NOTCH3 intracellular domain maintain a MES identity *in vivo* (32). In contrast, another study reported a spontaneous MES-to-NOR transition following engraftment of CD44-positive subpopulations isolated from SK-N-SH or IC-pPDXC-63 NB cell lines (33). Together, these findings suggest that maintenance or conversion of the MES state *in vivo* is highly context-dependent and influenced by intrinsic cellular properties, and potentially by microenvironmental cues.

Despite a shared MES background, tumorigenicity, phenotypic plasticity, and ALK signaling differed markedly across models and ALK variants. In GIMEN cells, which had limited tumorigenic capacity, only ALK^-F1174L^ and ALK^-R1275Q^ expression elicited tumor formation. Although ALK^-F1174L^ tumors exhibited higher ALK signaling activity assessed by ALK signature scores, they grew more slowly than ALK^-R1275Q^ tumors, and both tumor groups displayed distinct transcriptional profiles, highlighting differences in downstream signaling between these activating ALK variants.

The SH-EP model displayed a higher intrinsic tumorigenic potential relative to the GIMEN model, as all NSG mice implanted with transduced SH-EP cells developed tumors. However, each ALK variant drove distinct growth kinetics, phenotypic changes, and transcriptomic outcomes. Importantly, ALK-wt and ALK^-F1174L^ expression were associated with a partial reduction of MES identity and acquisition of neuroblastic and NOR features, as reflected by changes in tumor morphology, focal loss of CD44 expression in ALK^-F1174L^ and ALK-wt tumors, and the emergence of PHOX2B-positive regions in one third of ALK-wt tumors. This observation is particularly striking, as no specific driver of MES-to-NOR transition has been described to date, except with EZH2 inhibition (34). In contrast, the NOR-to-MES transition is better documented and has been described to be induced by PRRX1 expression, NOTCH3 signaling, or ARID1 depletion (20, 32, 34, 35). Since ALK is a component of the NOR signature (20), and its pharmacological inhibition has been reported to promote enrichment of MES-like cell populations, it remains important to determine whether, and to what extent, ALK signaling contributes to the maintenance of the NOR state in NB.

Conversely, ALK^-R1275Q^ expression in SH-EP tumors preserved a MES phenotype, while activating a distinct transcriptional program enriched in wound healing, extracellular matrix remodeling, and epithelial adhesion pathways, in accordance with the MES state. This included strong upregulation of hemidesmosome/desmosome-associated genes (*KRT14*, *KRT5, COL17A1, DSP, DSG2*), reflecting the stabilization of a highly adhesive MES state in SH-EP cells.

Furthermore, this transcriptional state aligns with metastatic behavior, as ALK-R1275Q tumors exhibited efficient dissemination, predominantly as numerous isolated cells or small clusters. These results are consistent with a previous study reporting enhanced dissemination in ALK-R1275Q/TH-MYCN-transgenic mice versus TH-MYCN, although the specific ECM and adhesion genes involved differed between the models (36). The upregulation of *KRT14* and adhesion complex genes in ALK^-R1275Q^ tumors may represent an alternative metastatic strategy involving collective migration and dissemination in the SH-EP model. Indeed, in breast cancer, KRT14-expressing cells are enriched in circulating tumor cell clusters and micrometastases, but are rare in established macrometastases, promoting collective invasion and early metastatic spread while maintaining intercellular adhesion (30). The role of KRT14 in the metastatic behavior of SH-EP-ALK^-R1275Q^ cells remained to be confirmed through functional studies.

Moreover, transcriptomic analyses of metastatic lesions further highlighted distinct adaptation strategies. ALK^-F1174L^ metastases showed limited transcriptional divergence from primary tumors, consistent with recent single-cell transcriptomic studies in NB showing that major transcriptional programs of NB tumor cells are largely preserved during metastatic progression (37, 38). In contrast, ALK^-R1275Q^ metastases exhibited greater transcriptional reprogramming, suggesting increased plasticity during dissemination and potentially early adaptation to the lung microenvironment. Together, these findings highlight how activating ALK variants determine distinct, context-dependent oncogenic programs that shape tumor identity, growth dynamics, and modes of dissemination in mesenchymal NB.

In NB patients, ALK mutations have been reported to be more frequent at relapse, either due to *de novo* emergence or increased variant allele frequency through clonal selection (39, 40). Clonal subpopulations carrying different ALK variants can expand or regress over time, reflecting both intrinsic differences in variant-dependent oncogenic potential and the influence of tumor genetic background (18, 41). ALK^-R1275Q^ was the most frequently detected ALK mutation in relapsed NB, showing significant enrichment compared to primary tumors, whereas ALK^-F1174L^mutant clones regressed in a subset of cases (39). In line with these observations, our *in vivo* data supports that ALK variants confer a context-dependent selective advantage. Specifically, ALK^-R1275Q^ promoted either tumor growth (GIMEN) or dissemination (SH-EP) while maintaining a MES transcriptional state associated with aggressive disease. In contrast, in GIMEN cells, ALK^-F1174L^ delayed tumor growth, consistent with the reported regression of ALK^-F1174L^ mutant clones in patients. Together, these findings highlight the heterogeneous and dynamic nature of ALK-driven tumor evolution and the role of cellular context in shaping clonal selection and disease progression.

Several limitations should be considered. Our study relies on cell line–derived xenograft models in immunocompromised mice, which do not fully recapitulate the native tumor microenvironment. Although the use of multiple well-characterized models allows assessment of ALK variant–specific effects, validation in patient-derived xenografts (PDX) models would be valuable to confirm the context-dependent signaling outputs of distinct ALK variants. Moreover, subcutaneous implantation may influence both tumor phenotypes and dissemination patterns, including the predominance of lung metastases. In addition, the SH-EP model displays epithelial features that may contribute to keratin gene expression, potentially reflecting model-specific characteristics limiting direct extrapolation to other NB contexts (42). Furthermore, bulk transcriptomic analyses do not capture intratumoral heterogeneity, and single-cell approaches will be required to better define the effect of specific ALK variants on cellular plasticity. Finally, cellular states were defined using *in vitro*-derived MES and NOR signatures only, without incorporating signatures from *in vivo* studies. Although suitable for the quite stable MES models studied here, this may not fully reflect the spectrum of cellular states present in patient tumors, where NOR cells predominate, and MES states are less clearly distinct. (20, 33, 43).

Overall, our study underscores that ALK-wt, ALK^-F1174L^, and ALK^-R1275Q^ drive distinct, context-dependent oncogenic programs, shaping tumor growth, cellular identity, and modes of dissemination. Although the precise molecular mechanisms linking individual signaling pathways to specific tumor behaviors remain to be elucidated, these findings highlight the importance of integrating mutation type, cellular identity, and tumor environment when considering ALK biology and therapeutic targeting.

## ACKNOWLEDGMENTS

We thank Dr. Carlo Fusco for providing the pLiVpuro_C vector. Histological analyses were performed at the Histology Facility (UNIL) and the EPFL Histology Core Facility with the precious help of Janine Horlbeck and Dr. Jessica Dessimoz, respectively. RNA sequencing analyses were performed at the iGE3 Genomics Platform of the University of Geneva (https://ige3.genomics.unige.ch). We also thank the Animal Facility (UNIL, Bugnon 27), the AGORA In vivo Center and In Vivo Imaging Facilities for their help with animal experimentation, as well as the Cardiology Assessment Facility (UNIL) for their support with echography in orthotopic experiments. Tissue slides were imaged with the instructions of the Cellular Imaging Facility (CIF, UNIL) using the Zeiss Axioscan Z.1 and Zeiss Axioscan 7 slide scanners. This work was supported by grants from the Swiss National Science Foundation (SNSF, # 310030_163407 and 310030_189074 to AMM), the FORCE Foundation (to AMM and RR), and the Stiftung Kinderkrebsforschung Schweiz (to AMM).

## AUTHOR CONTRIBUTIONS

MEN and LVS performed all major experimental work with the technical help of KBB. MEN, LVS and AMM designed the study and analyzed the data. NJ performed orthotopic cell implantations. NG and VP conducted bioinformatic analyses. NR provided help in pathological analyses of tumors and lung tissues. IJL, NR and RR provided resources. RR contributed to conceptual advice and fundings. MEN, LVS, NG and AMM prepared the figures, drafted and edited the manuscript. All authors read and approved the final manuscript.

## COMPETING INTERESTS

The authors declare no competing interests. N.R is an employee of Genentech since 7 February 2022.

## DATA AVAILABILITY

All data generated during this study are included in this article (and its Supplementary Information file). The RNAseq raw data generated in this study have been deposited at ENA (https://www.ebi.ac.uk/ena/browser/home). The processed RNAseq data have been deposited at ArrayExpress (https://www.ebi.ac.uk/biostudies/arrayexpress) under the accession E-MTAB-15992. Microscopy image datasets have been deposited in BioImage Archive (BIA, https://www.ebi.ac.uk/bioimage-archive/) under the accession number S-BIAD3866.

