## Supplementary information for "ALK Variants Differentially Modulate Neuroblastoma Tumor Behavior and Transcriptome in a Cellular Context-Dependent Manner"

This section includes the description of Supplementary Methods and Supplementary Figures and their Legends; and Supplementary Reference.

### Supplementary Methods

#### Immunocytochemistry

Immunocytochemistry (ICC) was performed on cytospin cell preparations. Cells were cytocentrifuged onto glass slides and fixed in Histofix (ROTI®Histofix, Roth AG) for 10 min at room temperature, followed by a 5-min wash in 1× TBS. Permeabilization was carried out for 10 min using Triton buffer (0.1% Triton X-100 Sigma-Aldrich, 0.05% $NaN_3$ in PBS), then washed three times in TBS. Anti-ALK (D5F3®, Cell Signaling 3633S) diluted 1:200 in TBS was applied overnight at 4 °C. After a 5-min TBS wash, slides were incubated with a anti-rabbit immunoglobulin–alkaline phosphatase secondary antibody (Abcam ab6729, 1:2500 in TBS) for 30 min at room temperature, followed by a 5-min TBS wash. Signal was developed using Liquid Permanent Red (Fast red substrate Kit, Abcam ab64254) for 10 min, then slides were rinsed in distilled water, counterstained with Hematoxylin Gill No.2 (Merk 105175) for 5 min, washed in distilled water, and mounted in glycerin. Staining was analyzed using the Olympus BX43 light microscope.

#### Immunoblotting

Cells and tumor xenografts were lysed in NP40 buffer: 50 mM Tris-HCl pH 8.0; 150 mM NaCl; 1% NP-40 and 1x protease inhibitor cocktail (Complete mini, EDTA-free, Roche, Mannheim, Germany) and centrifuged for 10 min at 15’000 rpm at 4°C. Protein concentrations were measured using Bradford method (Bio-Rad Laboratories, Hercules, CA, USA). Protein lysates were loaded on Mini-PROTEAN TGX Gels 4-15% 10 wells (Bio-Rad Laboratories), transferred to PVDF membrane (Bio-Rad Laboratories). Proteins were detected using monoclonal rabbit antibody for ALK (D5F3®, 1/1000, Cell signaling technologies, MA, USA) and monoclonal mouse antibody for β-Actin (1/10000, Sigma-Aldrich, MO, USA), then using secondary antibody goat anti-Rabbit IgG/HRP (1/10000, Dako, CA, USA) or peroxidase AffiniPure goat anti-mouse IgG (1/10000, Jackson ImmunoResearch, UK), respectively; and revealed using WesternBright Sirius Kit or WesternBright Quantum kit (Advansta, CA, USA). Membranes were imaged the Fusion FX6 EDGE multimodal imaging platform (Vilber Lourmat, Marne-la-Vallée, France).

#### Immunohistochemistry

Immunohistochemistry (IHC) was performed on paraffin-embedded blocks, using ALK monoclonal rabbit antibody (D5F3®, Cell signaling technologies, 1/250, Ag retrieval EDTA), CD44 (clone F.10.44.2[1], 1/100 Ag retrieval EDTA), PHOX2B (B-11, Santa Cruz, 1/200, Ag retrieval EDTA), KRT14 (LL002, Abcam, 1/200, Ag retrieval Citrate) and KRT5 (EPR1600Y, Abcam, 1/200, Ag retrieval Citrate). Sections were imaged using a BX43 Olympus light microscope and the CellSens Entry 1.18 imaging software (LEAD Technologies, Inc). Whole slides were scanned at the 10x or 20x magnification using the Zeiss Axioscan Z.1 or Zeiss Axioscan 7 instruments at the Cellular Imaging Facility (CIF, UNIL).

**Reverse transcription and real-time PCR**

cDNA was prepared from 0.5 µg of RNA using the PrimeScript™ reagent kit (TAKARA Bio Inc., Shiga, Japan). Real-time PCR was performed using the RotorGene 6000 real-time cycler (Corbett, QIAGEN). Cycling conditions were 95°C for 5 min, 40 cycles of 95°C for 10 sec, 60°C for 20s, and 72°C for 1s using the QuantiFast SYBR® green kit (QIAGEN). Relative expression levels were calculated using the ΔCt method with *HPRT1* and *SDHA* as housekeeping genes. Primers (Microsynth) are described in Supplementary Table 1.

| Gene | Gene description | Forward | Reverse |
|---|---|---|---|
| HPRT1 | Hypoxanthine Phosphori-bosyl transferase 1 | TGACACTGGCAAAACAATGCA | GGTCCTTTTCACCAGCAAGCT |
| SDHA | Human Succinate Dehydrogenase Complex Flavoprotein Subunit A | TGGGAACAAGAGGGCATCTG | CCACCACTGCATCAAATTCATG |
| m-SDHA | Mouse Succinate Dehydrogenase Complex Flavoprotein Subunit A | GCCAGGAACACTCCAAAAAC | ATTCATGATCCACCACTGGG |
| ALK | Anaplastic Lymphoma Kinase | TGTTGCCTCTCCTCGATGTG | TGTCCTCTCCGCTAATGGTG |
| KRT14 | Keratin 14 | AACGCCGACCTGGAAGTGAAG | TTGTCCACTGTGGCTGTGAGA |
| KRT5 | Keratin 5 | ATCGCCACTTACCGCAAGC | CCACTGCCATATCCAGAGGAA A |
| COL17A1 | Collagen 17A1 | CCAGGGTGAGAAAGGACCTC | TGTTCACCTCTTGGGCCT TG |
| DSP | Desmoplakin | TCA CCA GTG AAT GTT TGG GGT | GAT AGT CGC CGA TGG AGT TGT |
| ETV1 | E twenty-six variant 1 | CAG GAA CCC AGA GAT TTT GC | GGC CCT TTT CAA ACA TAC AGC |
| ETV4 | E twenty-six variant 4 | CAG CCA TGA ATT ACG ACA AGC | TCA CAC ACA AAC TTG TAC ACG |
| ETV5 | E twenty-six variant 5 | AGC TCT TCA GAA TCG TGA G | TCT CGA TCT GAG GAA TGC AG |
| DUSP6 | Dual-specificity phosphatase 6 | ACA GTG GTG CTC TAC GAC G | AAG ACA CCA CAG TTC TTG CC |
| SPRY2 | Sprouty RTK Signaling Antagonist 2 | TGC CTT GTT TAT GGT GTT ACC | GAG TTT TTA CAG CGG CAA CC |
| SPRY4 | Sprouty RTK Signaling Antagonist 4 | GGG TAC CTT TAA AGA AGA CCC | GGT GTT CTA CAC CTT CCC C |

**Supplementary Table 1:** List of primers used in the study.

### Supplementary Figures

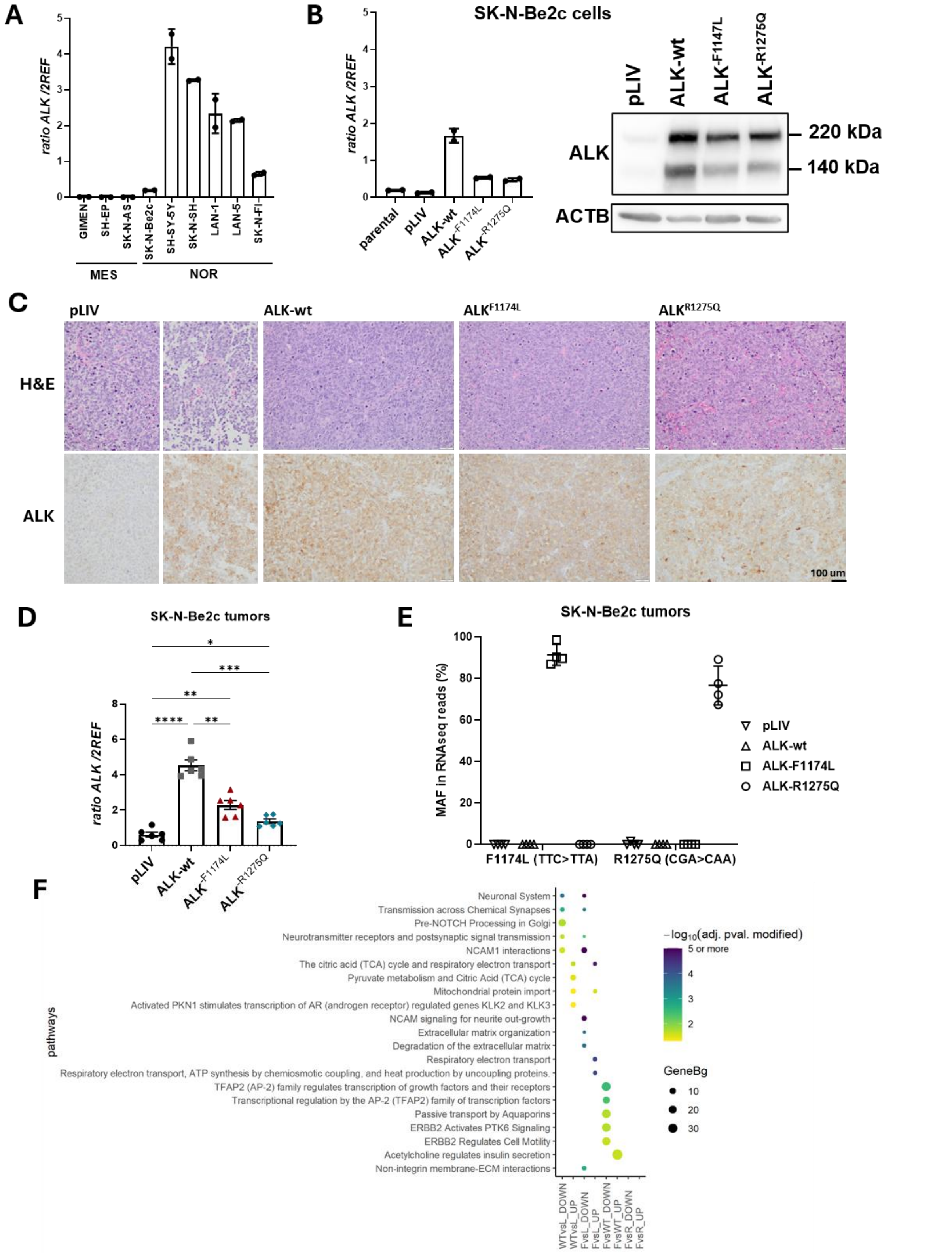


**Suppl. Figure 1. (A)** ALK mRNA expression ratios measured by RTqPCR in several MES and NOR NB cell lines in duplicates using *HPRT1* and *SDHA* as housekeeping genes. **(B)** ALK expression in SK-N-Be2c parental and transduced cell lines analyzed by RTqPCR as in (A) (left panel) and by immunoblotting (right panel). Actin-β (*ACTB*) was used as loading control. **(C)** Representative images of H&E and ALK expression detected by IHC in SK-N-Be2c tumors, scale bar = 100 µm. **(D)** ALK mRNA expression levels relative to *HPRT1* and *SDHA* control genes in SK-N-Be2c tumors analyzed by RTqPCR and plotted as individual values and mean ±SD. Brown-Forsythe and Welch ANOVA tests: * $p<0.05$, ** $p<0.005$, *** $p<0.0005$, **** $p<0.0001$. **(E)** Mutated allele fraction (MAF) of F1174L or R1275Q ALK alleles relative to wt sequence identified in RNAseq reads in each orthotopic tumor. Percentage of reads with the mutated allele vs total number of reads covering the sequence of the indicated hotspot mutation in each tumor are indicated in the graph with mean ±SD. **(F)** Dot plot of the top 5 (by adjusted p-values) enriched REACTOME pathways among upregulated (UP) and downregulated (DOWN) gene lists in each indicated comparison of SK-N-Be2c tumors. No pathways were enriched in ALK$^{-R1275Q}$ vs pLIV and ALK$^{-R1275Q}$ vs ALK-wt comparisons. Dot size represents the enrichment factor measured as Gene ratio/background ratio (GeneBg), and the color indicates the significance level (adjusted p-value, $-\log_{10}$ scale). L = pLIV, WT = ALK-wt, F = ALK$^{-F1174L}$, R = ALK$^{-R1275Q}$, UP = upregulated, DOWN = downregulated.

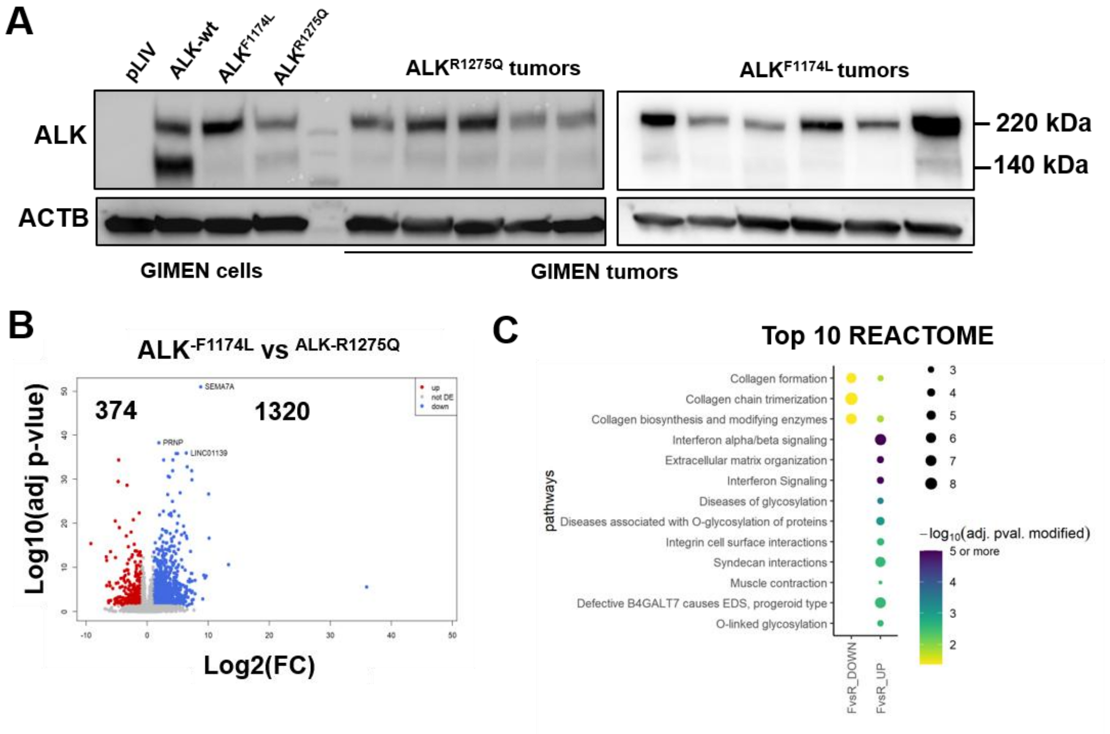


**Suppl. Figure 2. (A)** ALK protein expression analyzed by immunoblotting in transduced GIMEN cells and their derived tumors. ACTB was used as loading control. **(B)** Volcano plot representing the upregulated (blue) and downregulated (red) DE genes detected in GIMEN ALK$^{-F1174L}$ relative to ALK$^{-R1275Q}$ tumors plotted according to $\log_2$(FC) and $-\log_{10}$(adj pvalue). **(C)** Dot plot showing the top 10 REACTOME most significantly enriched among upregulated and downregulated genes in GIMEN tumors. Dot size represents the enrichment factor measured as Gene ratio/background ratio (GeneBg), and the color indicates the significance level (adjusted p-value, $-\log_{10}$ scale). F = ALK$^{-F1174L}$, R = ALK$^{-R1275Q}$.

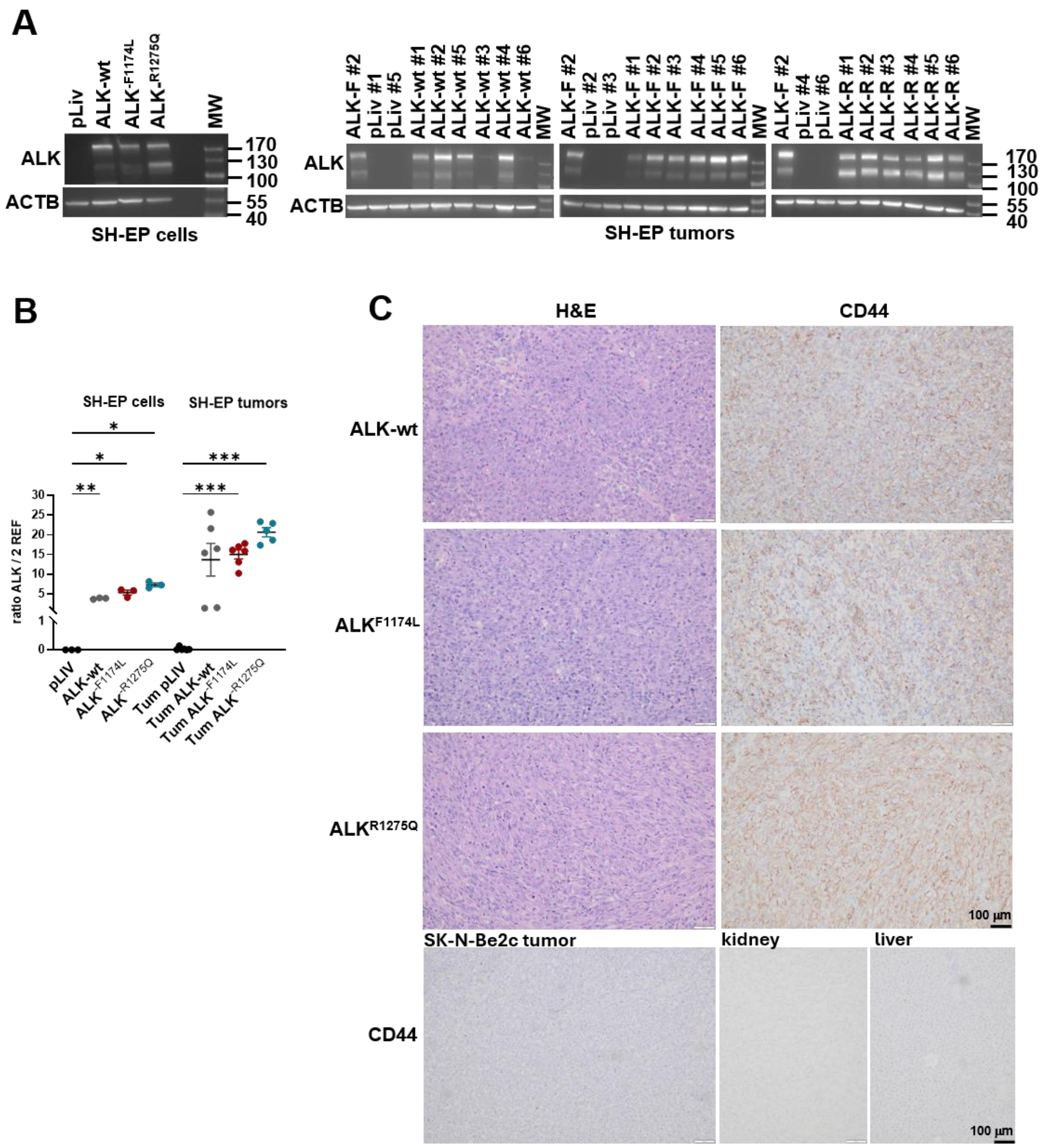


**Suppl. Figure 3. (A)** ALK protein expression analyzed by immunoblotting in transduced SH-EP cells and their derived tumors. ACTB was used as loading control. MW = molecular weights in kDa. **(B)** RTqPCR analysis of ALK mRNA expression levels relative to *HPRT1* and *SDHA* housekeeping genes measured in SH-EP transduced cells and their derived tumors. Individual values are plotted with mean ±SD. Brown-Forsythe and Welch ANOVA tests: * *p*<0.05, ***p*<0.005, ***p<0.0005. **(C)** Representative images of H&E staining and CD44 IHC in SH-EP-ALK derived tumors generated in athymic Swiss nude mice. Scale bare: 100 μm. Tumors developed in 3/6 mice in the ALK-wt group, 5/6 mice in the ALK-$^{F1174L}$ group, and 2/6 mice in the ALK-$^{R1275Q}$ group. A negative control panel represents the CD44 IHC staining on SK-N-Be2c control tumors (25), and on kidney and liver collected from NSG mice.

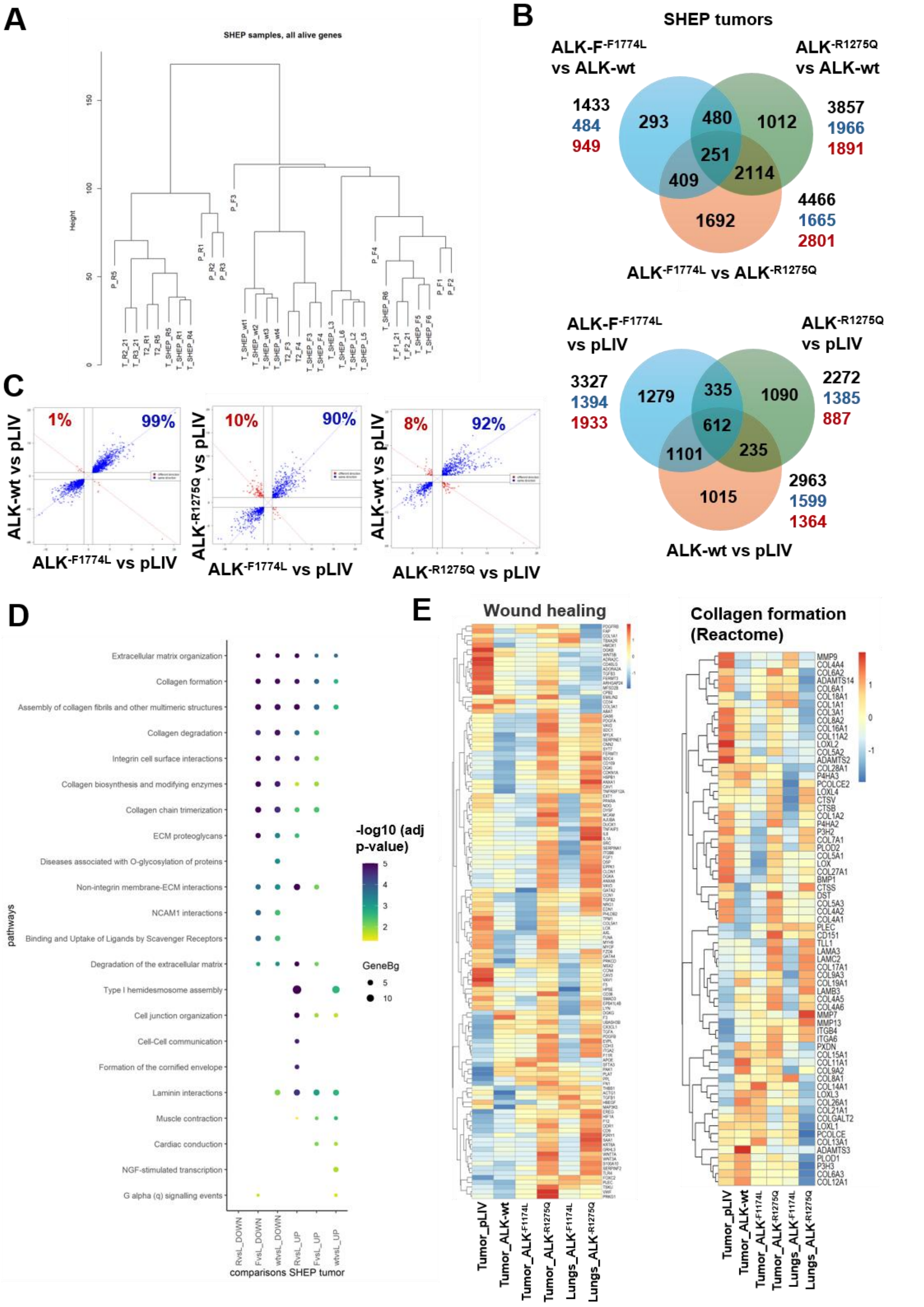


**Suppl. Figure 4. (A)** Unsupervised hierarchical clustering of SH-EP tumors and lungs samples. **(B)** Venn diagram showing the common and specific DEGs (adj p-value <0.05 et absolute log2FC ≥1) in SH-EP tumors vs control tumors pLIV (left panel) or vs ALK-expressing tumors (right panel). Total numbers of DEGs (black), upregulated (blue) and downregulated (red) genes are also indicated for each comparison. **(C)** Graphs showing the genes deregulated in common in the indicated comparisons with the log2(FC) and labelled in blue if deregulated in the same direction or in red if deregulated in opposite direction. **(D)** Dot plot showing the top 10 enriched REACTOME pathways sorted by adjusted p-values among upregulated and downregulated genes in the diverse comparisons of SH-EP tumors. Dot size represents the enrichment factor measured as Gene ratio/background ratio (GeneBg), and the color indicates the significance level (adjusted p-value, $-\log_{10}$ scale). **(E)** Heatmaps of the Wound healing (GO BP) and of the Collagen formation (REACTOME) pathways showing the DEGs of all group comparisons described in this study. Z-score scaled by row is shown for each sample group.

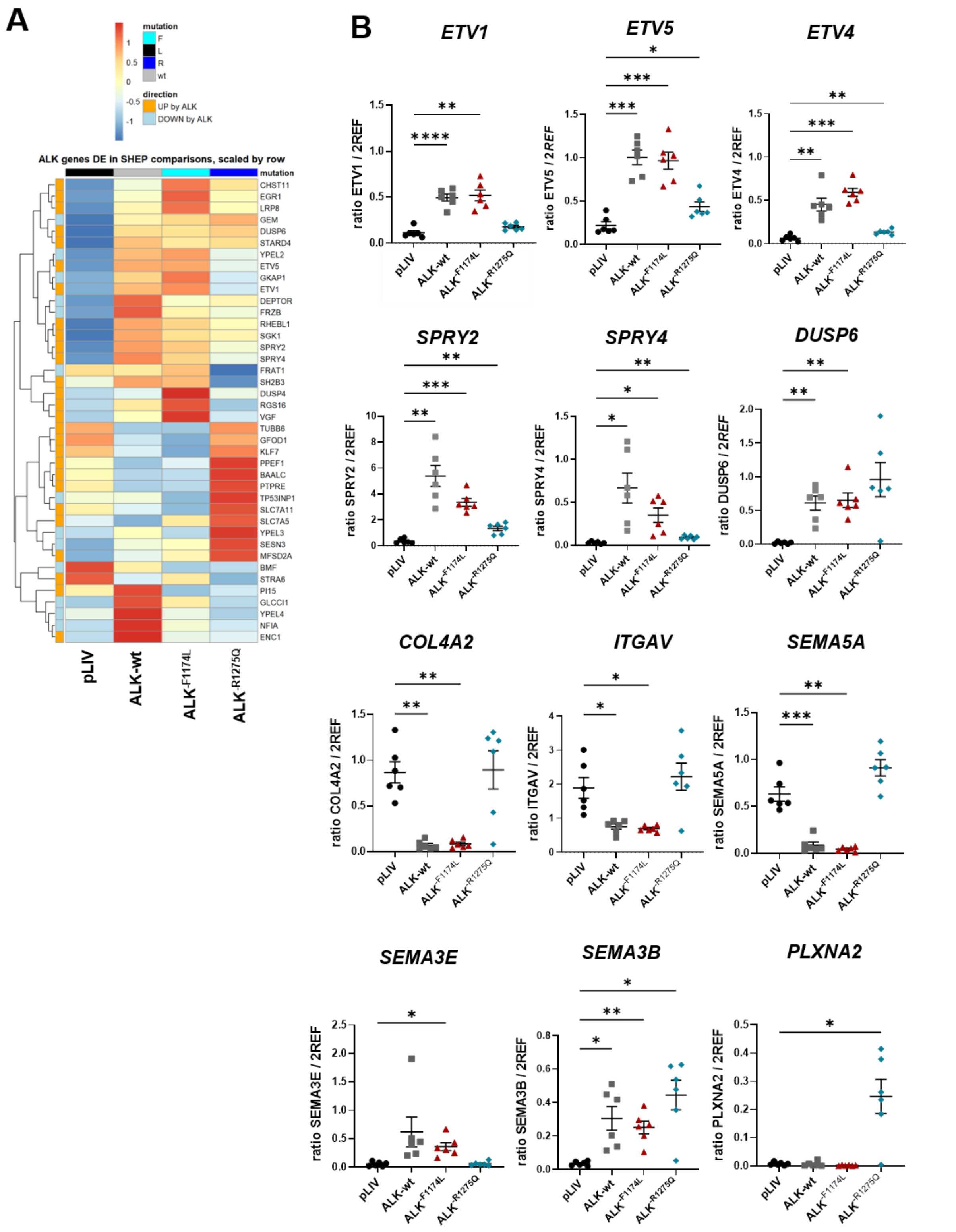


**Suppl. Figure 5. (A)** Heatmap of the ALK signature genes representing only the DEGs of the various comparisons between tumors. Z-score scaled by row are shown for each sample group. **(B)** mRNA expression ratios of genes of the ALK signature or selected genes, measured by RTqPCR in SH-EP tumors using *HPRT1* and *SDHA* as housekeeping genes. Values are plotted as individual values and mean ±SD. Brown-Forsythe and Welch ANOVA tests: * $p<0.05$, ** $p<0.005$, *** $p<0.0006$, **** $p<0.0001$.

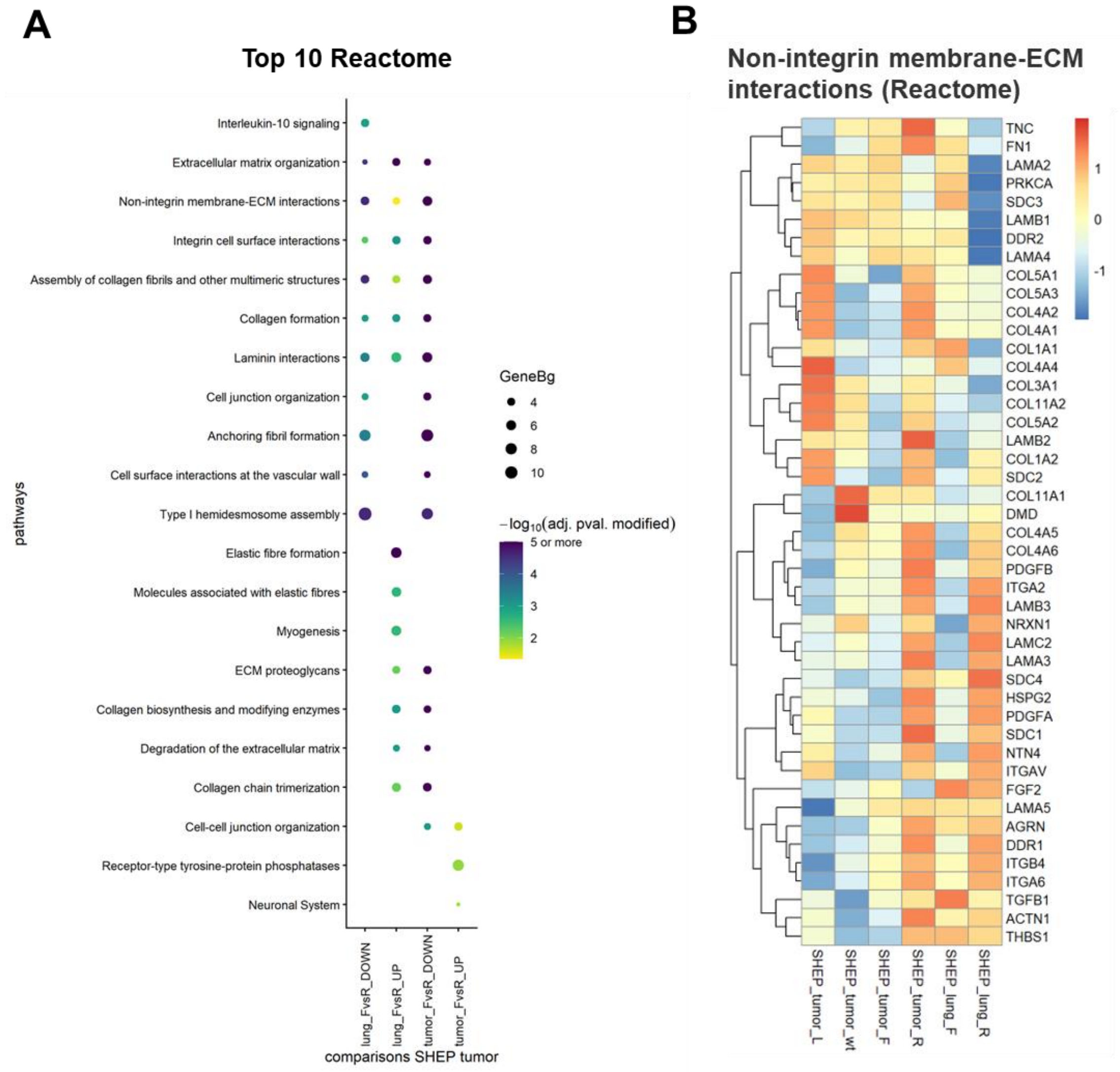


**Suppl. Figure 6. (A)** Dot plot showing the top 10 enriched REACTOME pathways sorted by adjusted p-values among upregulated and downregulated genes in ALK$^{-F1174L}$ vs ALK$^{-R1275Q}$ comparisons in lungs or in tumors. Dot size represents the enrichment factor measured as Gene ratio/background ratio (GeneBg), and the color indicates the significance level (adjusted p-value, $-\log_{10}$ scale). **(B)** Heatmap of the non-integrin membrane-ECM REACTOME pathway showing the DEGs of all group comparisons described in this study. Z-score scaled by row are shown for each sample group.

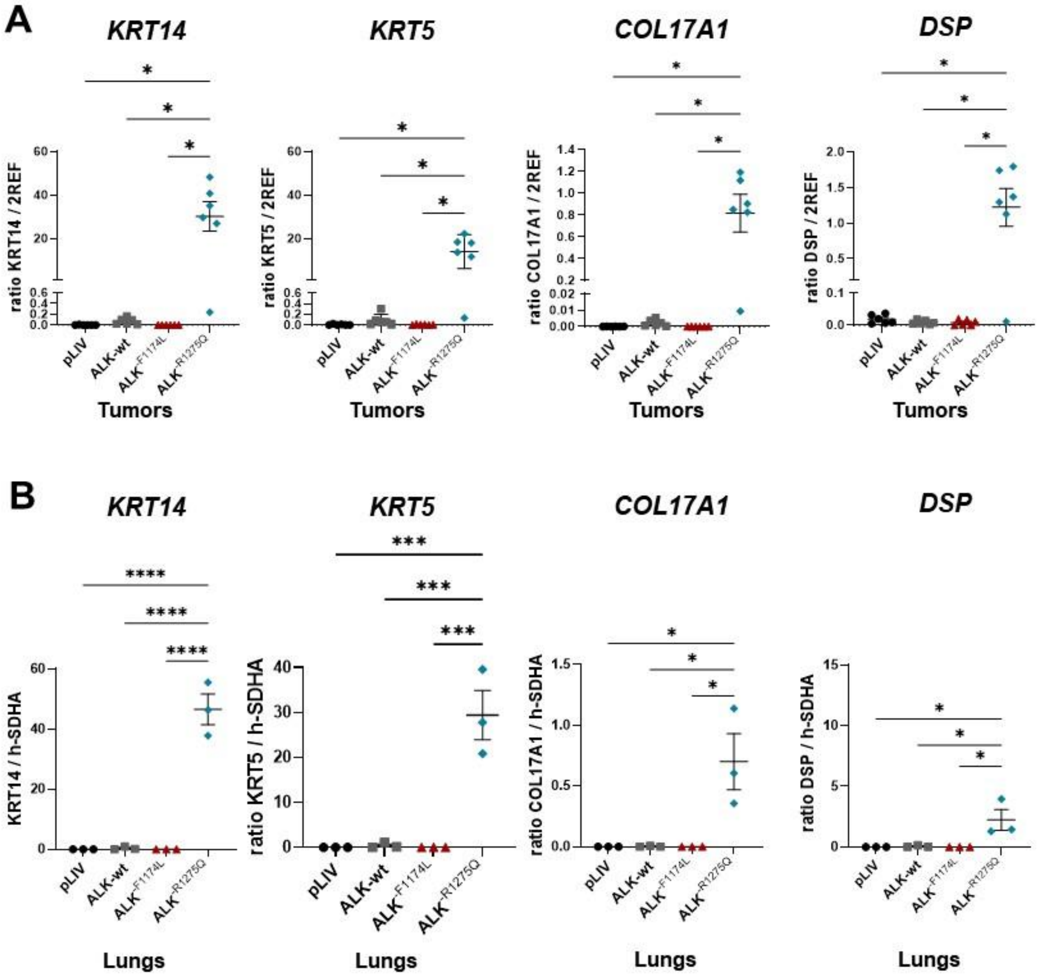


**Suppl. Figure 7. (A-B)** mRNA expression ratios of several hemidesmosome and desmosome genes measured by RTqPCR in SH-EP tumors using *HPRT1* and *SDHA* as housekeeping genes (A), or in SH-EP lung samples relative to *SDHA* expression (B). Values are plotted as individual values and mean ±SD. Brown-Forsythe and Welch ANOVA or Ordinary one-way ANOVA tests: * $p$<0.05, **$p$<0.005, ***$p$<0.0005, ****p<0.0001.
